# Chilean bee diversity: Contrasting patterns of species and phylogenetic turnover along a large-scale ecological gradient

**DOI:** 10.1101/2022.09.13.506767

**Authors:** Leon Marshall, John S. Ascher, Cristian Villagra, Amaury Beaugendre, Valentina Herrera, Patricia Henríquez-Piskulich, Alejandro Vera, Nicolas J. Vereecken

**Affiliations:** Agroecology Lab, Université libre de Bruxelles (ULB), Boulevard du Triomphe CP 264/2, B 1050 Brussels, Belgium; Naturalis Biodiversity Center, Darwinweg 2, 2333 CR Leiden, The Netherlands; Department of Biological Sciences, National University of Singapore, 14 Science Drive 4, Singapore 117543, Singapore; Instituto de Entomología, Universidad Metropolitana de Ciencias de la Educación, Santiago, Región Metropolitana, Chile; Departamento de Biología, Universidad Metropolitana de Ciencias de la Educación, Santiago, Región Metropolitana, Chile

**Keywords:** Generalized dissimilarity modelling, phylogenetic diversity, climate, land-use, β-diversity, solitary bees, turnover, nestedness, global biodiversity hotspot, ecoregions, endemism, biogeography

## Abstract

Chile’s isolation and varied climates have driven the evolution of a unique biodiversity with a high degree of endemism. The Mediterranean-type biome of Central Chile is one of 35 global biodiversity hotspots and has been highlighted as one of Chile’s most endangered areas. It is threatened by anthropogenic land use change impacting the integrity of local biomes and associated species. This area holds the most extensive collections of the country with high endemicity regarding native bee species. Characterising habitat requirements of bees is a pressing priority to safeguard them and the ecosystem services they provide.

We investigated broad-scale patterns of bee diversity using newly accessible expert-validated datasets comprising digitized specimen records from both Chilean and US collections and novel, expert-validated type specimen data for the bees of Chile. We used a generalised dissimilarity modelling (GDM) approach to explore both compositional and phylogenetic β-diversity patterns across latitudinal, altitudinal, climate and habitat gradients in well-sampled bee assemblages in Central Chile. Using the GDM measures of increasing compositional and environmental dissimilarity we categorised and compared the most important drivers of these patterns and used them to classify ‘wild bee ecoregions’ (WBE) representing unique assemblages.

Turnover of bee assemblages was explained primarily by latitudinal variation (proxy for climate) along Chile. However, temperature variations, precipitation and the presence of bare soil also significantly explained the observed patterns. In comparison, we observed less turnover in phylogenetic biodiversity corresponding to spatial gradients. We were able to develop six *de novo* ecoregions (WBE) all with distinct taxa, endemic lineages, and representative species. The WBE represent distinct spatial classifications but have similarities to existing biogeographical classifications, ecosystems and bioclimatic zones.

This approach establishes the baseline needed to prioritise bee species conservation efforts across this global biodiversity hotspot. We discuss the novelty of this classification considering previous biogeographical characterisations and its relevance for assessing conservation priorities for bee conservation. We argue that Chile’s WBE show areas requiring funding for bee species exploration and description, distribution mapping and strengthening of conservation policies.

## Introduction

Geographic isolation and size of area mediate patterns of biodiversity by limiting the size and exchange of populations and extinction rates (MacArthur & Wilson, 1967). This in turn drives differences in compositional and phylogenetic diversity (Whittaker & Fernández-Palacios, 2007). South America has some of the most biodiverse regions in the world (Ulloa Ulloa et al., 2017), due to complex differences in abiotic and biotic factors across the many habitat types, which drives geographic isolation (Antonelli & Sanmartín 2011; Hughes et al., 2011). Biodiversity in South America has also been enriched by cycles of interchange between regions and biomes (Antonelli et al., 2018; Williams et al., 2022). Interestingly, Chile has a moderate species richness, lower than Amazonia, the Northern Andes and other regions of South America with globally maximal diversity for many groups but has extraordinary endemism. Chile is a good example of the effects of geographic isolation on biodiversity as it has many natural barriers to dispersal. The country is bordered by the Pacific Ocean along its western and southern coasts, by the high Andes Mountain range in the east, and by the Atacama Desert in the north (Veblen et al., 2007). Combined with significant long-term climate fluctuations, its geographical isolation has resulted in a unique biodiversity with high rates of endemism across many groups of animals and plants (Rovira et al., 2008). Within its territory, the Mediterranean-type biome of Central Chile is considered as one of 35 global biodiversity hotspots (Armesto et al., 2007; Myers et al., 2000; Moreira-Muñoz, 2011) and often referred to as a threatened region of utmost conservation priority (Mittermeier et al., 2004).

Nature’s contributions to people (NCP) (Díaz et al., 2018), has experienced an increasing deterioration in Central Chile through deforestation, intensive industries (*e.g.*, agriculture and mining), pollution and the spread of exotic species and its pathogens (Freitas et al., 2009; Underwood et al., 2009; Schulz et al., 2010; Carranza et al., 2020; Basualdo et al., 2022). Historically, there has also been a lack of financial support of the Chilean State for biodiversity conservation (Waldron et al., 2013). As a consequence, underregulated changes in land use due to industrialized forestry and agricultural activities have been identified as among the major threats to this hotspot’s wildlife (Hernández et al., 2016). This has significantly compromised the integrity of natural and semi-natural habitats and their biodiversity (Alaniz Baeza et al., 2016).

As has been also demonstrated worldwide (Potts et al., 2016), wild bees in Chile are a conservation concern due to their irreplaceable roles within ecosystems (Henríquez-Piskulich et al., 2021; Vieli et al., 2021). The health, survival and associated ecosystem services provisioning of native bee assemblages are increasingly jeopardized through the decline of their floral hosts and loss of suitable nesting resources as habitats become fragmented due to major shifts in land use (Potts et al., 2010). In Chile, researchers have documented the faunistic uniqueness and high endemism of Chile’s bee fauna (Moldenke 1976b; Packer and Ruz 2016; López-Aliste et al., 2021). Several authors express serious concerns about the threats faced by some of its most iconic Apoidea species, such as the world’s largest bumblebee *Bombus dahlbomii* Guérin-Méneville, 1835 (Hymenoptera: Apidae) (Morales et al., 2013). Considering the geographic uniqueness of Chile’s territory and its potential effects on wild bee species distribution, the situation of several Chilean native bee lineages may be even worse than what it is been reported for the popular *B. dahlbomii.* Despite the current conservation status, the giant bumble bee still presents a relatively broad distribution in comparison with several less attended wild bee lineages found in Chile (Morales et al., 2022). Many native bee species in Chile, with specific ecological needs, are endemic and restricted to a given region in Chile’s hotspot (Orr et al., 2021a). Some are highly dependent on specific floral resources, such as the solitary oil-collecting genera *Centris* Fabricius 1804 and *Chalepogenus* Holmberg 1903 (Hymenoptera: Apidae), both with species associated to Calceolariaceae with an array of species found at restricted locations of central and northern Chile (Murúa, 2020). Thus, it is possible to hypothesize that the patterns and drivers of wild Apoidea species diversity in Chile may reveal multiple and different conservation priorities if the unique characteristics of Chile’s biogeography are considered.

Given the potential conservation threats currently active in Chile, it is urgent to determine the patterns and drivers of bee compositional and phylogenetic diversity and identify bee-specific priority areas as part of a scientifically informed management strategy to preserve biodiversity for nature and people (Freitas et al., 2009; Gray et al., 2016; Martinez-Harms et al., 2021). Ultimately, such endeavours should include native bee diversity and endemism to strengthen the coverage, quality and connectivity of protected areas and improve ecosystem representation (Aycrigg et al., 2013; Rodrigues et al., 2004). The application of biogeographical knowledge on bees may help promote the use of biodiversity-friendly land management. Which will not only protect native bee species but will allow us to keep benefiting from the ecosystem services they provide.

In this study, we aimed to identify large-scale environmental drivers of changes in Chile’s wild bee assemblages (i.e., β-diversity), both in terms of species composition and shared evolutionary history (phylogenetic diversity), as well as to demarcate and characterize natural clusters or ‘wild bee ecoregions’ (WBE) and their contribution to the conservation of Chile’s unique wild bee diversity, while also comparing and contrasting them with historical biogeographic classifications of Chile (Kuschel,1960; Morrone, 2006, 2015; Peña 1966). Specifically, we compiled the largest-ever database of occurrence records relevant to the bees of Chile, including digitized museum specimen records and citizen science databases collected between 1851 and 2022, altogether comprising 59,000 occurrences for 444 species of Chilean bee species. We used national sources as well as the extensively digitized collections of the inventorying expeditions from the long-term collaboration between the US and Chile, stored at the American Museum of Natural History (AMNH). Using the validated dataset, we addressed the following three key questions: (i) What is the temporal and spatial distribution of species occurrence data and how does this differ between the main data sources? (ii) What are the large-scale distribution patterns of taxonomic and phylogenetic diversity patterns of wild bees in Chile, and what are the environmental drivers underlying these patterns? (iii) Can we classify *de novo* ecoregions of wild bee diversity (WBE) using their occurrences and environmental conditions and are these WBE consistent with biogeographical regionalisations based on other organisms?

## Methods

### Study area and species data

We conducted our study in the biodiversity hotspot that is found in Central Chile (Mittermeier et al., 2004). We defined Central Chile according to administrative regional boundaries the area from Región de Los Ríos in the south to the Región de Atacama in the north (Armesto et al., 2007), which contained most species occurrence data (>90% of raw occurrence records). The far North of Chile, north of the study area delimited here, is excluded due to an extreme difference in climatic conditions leading to a highly divergent bee fauna, much more limited data availability (the largest collection for this region has not yet digitized) and a lack of access to recent collections. The far South is excluded because it is too cold and wet to support a rich bee fauna (Michener, 1979; Orr et al., 2021a). Works by Packer et al., (e.g. Dumesh & Packer 2013; Packer & Graham, 2020), have documented a rich bee fauna in Northern Chile (i.e. Arica and Parinacota that was less well surveyed historically). At present, over 470 species have been described for Chile (Ascher and Pickering, 2022), many well-curated morphospecies await description, and more remain to be collected and recognized as additional areas, underrepresented in current collections, are surveyed and integrative approaches to taxonomy are applied (Packer & Ruz 2016). A handful of researchers and research groups have produced most species descriptions, much of which occurred in the 70s and 80s with the work of Toro and his collaborators (*e.g.,* Ruz & Toro 1983, Toro & Moldenke 1979). More recently, contributions have been made through classical taxonomy as well as modern integrative taxonomy approaches (*e.g.,* Ferrari 2017, Monckton 2016, Vivallo 2013).

There exists well-curated and serviceable digital specimen data for wild bees, in both Chile and the American Museum of Natural History (AMNH) in New York. We collated species occurrence records from multiple sources. We obtained digitized museum records from AMNH (Ascher, 2016), which included 28,779 specimens of 255 species, vouchers from a series of expeditions by Jerome G. Rozen, Jr., and Chilean colleagues investigating the life history of bees (e.g., Rozen 1970, Rozen & Ruz 1995). Bees digitized at the AMNH also include specimens collected by well-known Chilean entomologists such as L. E. Peña, H. Toro (synoptic collection), M. A. Fritz, and A. Ugarte-Peña. We downloaded digitized museum records from the Pontificia Universidad Católica de Valparaíso (PUCV) via GBIF (López-Aliste et al., 2021; https://doi.org/10.15468/6knwyq). This dataset included 35,743 records from 168 species and including extensive collections by H. Toro, L. Ruz and their students. We supplemented these two datasets with information from three other sources: (i) data from type specimens of taxa described from Chile (Ascher, unpublished). (ii) iNaturalist research grade observations of wild bees from Chile downloaded via GBIF (Occdownload Gbif.org 11 February, 2022; https://doi.org/10.15468/dl.9z642v) including 1,829 observations of 88 species mostly identified or validated by JSA; (iii) a dataset of 1,760 wild bee occurrences from 90 species collected by authors CV, PH and AB, Instituto de Entomología, Universidad Metropolitana de Ciencias de la Educación in 2016, 2017 and 2018 from (a) Farellones, a mountain village in the Lo Barnechea municipality of Santiago Province, (b) within the City of Santiago, (c) west of the Curacaví commune in the Melipilla Province of central Chile’s Santiago Metropolitan Region, (d) the Quebrada de Córdova wetland, in the Valparaíso region and (e) from Los Molles and north of Los Molles in the Valparaíso and Coquimbo regions.

Citizen science and museum collection data contain both spontaneous observations and those resulting from systematic sampling, which represent a mix of spatiotemporally unique occurrences and those comprising (by design or fortuitously) records from many years for a single locality. Therefore, we first cleaned this data collection by (i) using a checklist of the wild bees of Chile (source files for Ascher and Pickering, 2022); to check all records for synonyms and update incorrect records to the currently accepted binomial name. (ii) We checked the location information against coordinates and removed records where the two did not match. (iii) We compared the distribution of each species to its known regional distribution and noted discrepancies and determined those which were correct and those which were errors (iv) We aggregated all records where the date, location and species names where the same. (v) Finally, we excluded honey bees, *Apis mellifera* L., 1758, from the dataset as this represents an exotic species that is not sampled by melittologists in a way comparable to other bees.

To conduct generalized dissimilarity modelling (GDM) and compare within region diversity metrics, the dataset should have both presence and a reliable estimate of absence data. We calculated β-diversity metrics only at sites where considerable sampling had occurred. To deal with any inherent biases in collection, we created a set of three rules to define and only work with well-sampled localities within our chosen study area of Central Chile:

1. Sites had to be visited in at least three different years to ensure that the sites represented repeated visits.
2. Sites had to be visited in at least two different months of the year, to improve the coverage of within year variability of flying periods of different species.
3. Sites needed to have recorded occurrences of at least 10 different species, as to remove sites that might have represented limited collections and single species studies.

Additionally, based on expert opinion and historical treatments, we removed certain genera (especially of family Halictidae) from the GDM analysis, which we expected to be poorly determined (*Lasioglossum*, *Halictillus*) or subject to blatant digitization biases such as incomplete data entry of curated specimens (*Ruizantheda*). To account for variation in the resolution of the coordinates of the different occurrence records over time, we aggregated all records to a 5×5 km grid cell (hereafter referred to as a ‘site’). This allowed us to aggregate the well-sampled sites together with the extra specimens collected at the same location. Therefore, there are two separate datasets that classify the wild bee assemblages of the Central Chilean study area that are used throughout the analysis. Firstly, the full cleaned, dataset of temporally and spatially unique occurrences (hereafter ‘unique bee records’) found in Central Chile. This dataset is used to characterise the data and analyse the wild bee assemblages found in WBE *a posteriori*. The second dataset is a subset of the unique bee records with only those 5×5km cells classified as well sampled (hereafter ‘well sampled assemblages’). This dataset is used to train the GDMs and is the basis for the WBE classifications. We use ‘assemblages’ to refer to all the species that occur in in a specific place (e.g., WBE or well-sampled grid 5×5 km cell).

### Wild Bee Phylogeny

We calculated phylogenetic dissimilarity using two methods. First, a molecular phylogeny was produced based on all publicly available sequences of the mitochondrial gene cytochrome oxidase I (COI) available on GenBank (Benson, et al., 2014). All COI bee sequences were aligned using the MUSCLE multiple alignment algorithm in the msa package (v1.22.0; Bodenhofer et al., 2015). The alignment was converted into a pairwise distance matrix using dist.ml in the Phangorn package (v2.5.5; Schliep 2011). This distance matrix was converted into a tree using neighbor-joining tree estimation (Saitou & Nei, 1987). The Generalized Time-Reversible model with invariable sites and gamma model of rate heterogeneity (GTR + I + Γ) was selected as best nucleotide substitution model for our aligned COI sequences (Tavaré 1986), and a maximum likelihood phylogenetic reconstruction was made optimizing the model parameters (Figure S2). To account for the large number of missing species when using molecular data, we also calculated phylogenetic dissimilarity using a tree based on the taxonomic hierarchy of Linnean classification (Rank, Family, Subfamily, Tribe, Subtribe, Genus, Subgenus, Species), which has been shown to produce comparable results to a molecular phylogeny for Belgian bees (Vereecken et al., 2021), using the ‘ape’ package (v5.3) (Paradis and Schliep 2019). We built a polytomous, ultra-metric tree where all bee species recorded were grouped together according to their taxonomic classification (Michener 2000, Ascher and Pickering 2020, Ascher, unpublished). Branch lengths were computed using the ‘compute.brlen’ function from the ‘ape’ package. To compute phylogenetic beta diversity metrics at the site level, we used both the trees, separately, and the community matrix of well-sampled assembalges (presence/absence of species per site) as inputs of the ‘phylo.beta.pair’ function from the ‘betapart’ package (version 1.5.1) (Baselga et al., 2020). We extracted the dissimilarity matrix based on the Sorensen derived pair-wise phylogenetic dissimilarity. This matrix was used as the input for GDM.

### Environmental data

For the GDM analysis, we started out with an extensive set of ecologically relevant environmental variables to assess how they affect species compositional turnover. Climate variables included the global maps of the 19 Bioclims, as well as minimum, maximum and mean values of solar radiation and wind speed (v2.0 available at http://www.worldclim.org/ (Fick and Hijmans 2017, Hijmans et al., 2005). These were downloaded at 30 arc seconds resolution. Soil variables were extracted from the global WISE derived soil properties database also at 30 arc seconds resolution (available on the ISRIC Soil Data Hub; http://isric.org/explore/isric-soil-data-hub). The variables extracted included sand, silt and clay proportions, volume of coarse fragments, saturated water content and bulk density of the soil as these variables are likely to impact the potential habitat suitability for various wild bee species, especially ground-nesting species, which have consistent preferences for excavating nests (Cane 1991). Global land cover was obtained from Copernicus global land service at a 100×100 m resolution (downloaded in May 2019 from https://land.copernicus.eu/global/products/lc) (Buchhorn et al., 2020). We extracted bare/sparse vegetation, cropland, herbaceous grassland, shrubland, trees, permanent snow & ice, built-up areas. Finally, we included elevation at 3 arc seconds resolution (available at http://www.marine-geo.org) (Ryan et al., 2009). Each set of variables was cropped and masked to the extent and outline of our Central Chile study area using the raster package in RStatistics (v3.1.5) (Hijmans 2020). To match the resolution of the sites, each variable was then aggregated to a 5×5 km resolution (grid cell) to match the species data. For the climate, soil and elevation data we took the mean value of all the cells within each 5×5km grid cell and for the land cover data we took the percentage cover of each class within the grid cell. Therefore, before beginning the analysis we had an available set of 40 variables (Figure S1).

### Generalized dissimilarity modelling

To quantify the geographic and environmental patterns of wild bee species compositional turnover in Central Chile, we used GDM. All GDM analysis were conducted in R using the GDM package (v1.4.2) (Fitzpatrick et al., 2020). Generalized dissimilarity models are an extension of matrix regression and aim to model compositional dissimilarity between sites as a response to a set of explanatory environmental variables. These models represent an improvement on certain linear regression approaches by accounting for two common types of non-linearity found in ecological data (Ferrier et al., 2007). Observed dissimilarity and the rate of dissimilarity usually present a non-linear relationship along environmental gradients. When fitting a GDM, predictor variables are standardised, realised as I-spline basis functions, and fitted with a maximum-likelihood estimation (Ferrier et al., 2007). The height and the slope of the I-spline for each variable explains the amount and rate of the compositional turnover (Ferrier et al., 2007, Fitzpatrick et al., 2013). To select the suitable variables for a GDM we follow the procedure proposed by Williams *et al.,* (2012). For each pair of variables with a Pearson’s correlation coefficient greater than 0.7 we fitted a single GDM model and compared the percentage deviance explained, selecting the variable with the highest value. These variables were included together with all other non-correlated variables in a full-model GDM (see Figure S1). We used a backwards elimination model selection procedure. Each step removed the variable with the lowest coefficient until the final model was selected as the first model where all variables were significant at a 90% confidence (p-value < 0.1). The selected variables were then used to fit a GDM. We fitted a GDM with and without geographic distance and compared the percent deviance. We used Bray-Curtis distance to measure dissimilarity in species composition, as it gives more weight to double occurrences (Legendre and Legendre 2012) and should therefore reduce the biases resulting from false absences. We also weighted the sites based on species richness. This was done to further decrease the effects of different sampling artefacts in the data, given that richer sites were more intensively sampled. Three I-spline functions were used for all variables. A greater number of I-splines (five and seven) were tested, but these did not significantly change the results and the final models used three. The same steps were then repeated using phylogenetic distances between sites with a Sorensen derived pair-wise phylogenetic dissimilarity matrix as the response variable. The performance of the compositional GDM used to make predictions was assessed using cross-validation, we observed how well the model predicted the dissimilarity within a randomly withheld 20% of sites. This process was repeated a thousand times and the model performance metrics, including mean absolute error, where averaged (Mokany et al., 2022).

### Analysis of assemblage patterns

The resulting GDM was used to project compositional dissimilarity within the Central Chile study area. The final explanatory factors included in the best model were converted to their corresponding I-spline function values, which can be interpreted as a measure of their predicted biological importance to either species composition or PD. These transformed explanatory variables were then used to fit a principal component analysis (PCA). The coordinates from the first three axes of the PCA were then fitted to the entire region for each 5×5 km grid cell, resulting in three raster objects reflecting each axis of the PCA. The three uncorrelated raster layers were assigned to an RGB colour palette to show areas of increasing compositional dissimilarity and to be used for clustering regions with unique wild bee compositions (Hargrove and Hoffman 2004, Ferrier et al., 2007). Using k-means clustering (with max 10,000 iterations), we classified Central Chile into clusters based on predicted ecological dissimilarity. We chose the number of clusters based on the average of multiple indices with the NbClust package in R statistics, using a starting point of five clusters (where 80% of variance was explained) (v3.0) (Charrad et al., 2014). We took the mean of six indices after removing the minimum and maximum cluster values. The indices used were “ch”, “kl”, “hartigan”, “silhouette”, “gap”, and “ratkowsky” (Krzanowski and Lai 1988, Caliński and Harabasz 1974, Hartigan 1975, Rousseeuw 1987, Ratkowsky and Lance 1978, Tibshirani et al., 2001). The resulting clusters of compositional dissimilarity are henceforth referred to as “wild bee ecoregions” (WBE). The resulting WBE are then discussed in relation to previous historical, biogeographic regionalisations of Chile (see Morrone 2015).

For each bee ecoregion, we calculated four measures of phylogenetic biodiversity at both the site and bee ecoregion scale, and we used both; (i) the tree based on the molecular phylogeny, which only included species for which we had a COI sequence but more accurately represented branch lengths and the (ii) taxonomic hierarchy tree, which included all species but did not have accurate measures of branch length, resulting in 16 different values for phylogenetic biodiversity. We measured, (i) Phylogenetic species variability (PSV; the extent of relatedness between species in a community), (ii) Phylogenetic species richness (PSR; PSV multiplied by the species richness), (iii) Phylogenetic species clustering (PSC; a measure of the distance between branch tips across a community phylogeny) and (iv) phylogenetic diversity (PD; the sum of the total branch length) (Faith 1992; Helmus et al., 2007) using the picante package in R (v1.8.2) (Kembel et al., 2010). For the site level analysis, we took all 5 x 5km grid cells with at least three species records, to expand the breadth of the assemblages in each region, for the bee ecoregions we used all species records from the full database. At the site scale we also calculated compositional β-diversity by re-sampling 10 pair-wise site comparisons 100 times per region using Sorenson dissimilarity and splitting into the turnover and nestedness components, using the beta part R package (v1.5.2) (Baselga et al., 2020). Finally, we compared both phylogenetic β-diversity and β-diversity between regions using ANOVA.

### Endemism & Indicator Species

For each bee ecoregion we classified the bee species and species pairs that were most indicative of these assemblages using two datasets, all unique bee records and the well-sampled assemblages. All analyses of indicator species were conducted with the indicspecies package in r (version 1.7.9) (De Caceres and Legendre 2009). Species indicator value was calculated based on two criteria: (1) The probability, given that the species has been observed, that the surveyed site is part of the target site group; and (2) the probability of finding the species in sites that are part of the target site group. All calculations of the relationship between species and sites (Pearson’s phi coefficient of association) were corrected based on the number of sites in each site grouping. The bee species that demonstrated statistically clear (p<0.05) indicator values were reported as indicator species for each site group (bee ecoregion). The indicator species with occurrence wholly nested within other species occurrence were removed.

## Results

### Characterising species data

The cleaned dataset of 60,501records included 11,664 temporally and spatially unique occurrences (hereafter ‘unique bee records’) of 423 bee species (approximately 87% of species known for Chile), collected between 1851 and 2022 (Figure 1). During cleaning, a total of 369 unique bee records were removed (from PUCV and iNaturalist datasets) as the species mapped to outside its confirmed regional range. Across the five bee families found in Chile there were 3,798 unique bee records bee records of 82 species of Apidae, Colletidae (2,832 of151), Megachilidae (2,406 of 68), Halictidae (1,347of 53), and Andrenidae (1,281 of 69). There were 128 species recorded only once in the full dataset and 29 recorded twice. Ninety-eight percent of the occurrences were recorded post-1950 and 22% of the occurrences post-2000. The full database of digitized records shows considerable evidence of uneven sampling over time before 1950 (R^2^ = 0.36) based on the relationship between a species’ number of records and its area of occupancy. The best-sampled period was between 1970 and 2000 (0.85), representing the peak collecting periods of Peña, Rozen, and Toro, before decreasing again in recent decades (0. 40). Across all years, Colletidae showed the most even sampling (0.87). These results support the decision to aggregate all data together for the β-diversity analyses. The number of occurrences varied significantly between political regions, but unique bee records were recorded from all of Chile’s 16 regions. The highest number of species was recorded from Coquimbo with 1,835 unique bee records of 164 species, and the highest number of unique bee records from the Región Metropolitana de Santiago with 2,604 unique bee records of 149 species. The lowest number of unique bee records and species belong to the two most southern regions, Magallanes with 33 unique bee records of nine species, and Aysén with 47 unique bee records of 20 species.

**Figure 1.**
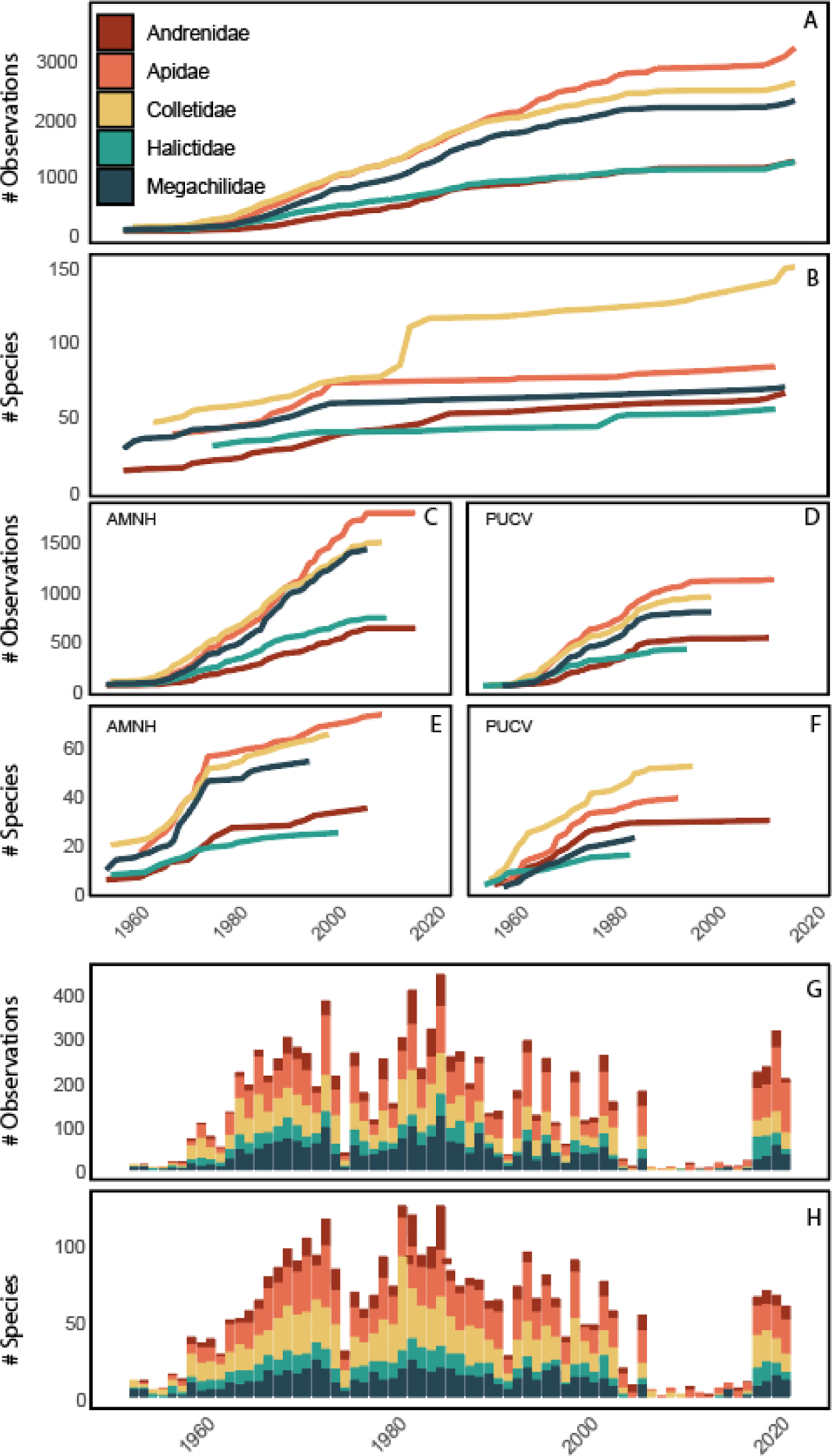
Summary of number of unique bee records and wild bee species recorded in the full dataset without duplicate date and location records. (A) Cumulative number of unique bee records recorded over time for each of the five bee families found in Chile. (B) Cumulative number of species recorded over time for each of the five bee families found in Chile. (C) Cumulative number of unique bee records recorded over time for each of the five bee families found in Chile based on the dataset obtained from the American Museum of Natural History (AMNH). (D) Cumulative number of unique bee records recorded over time for each of the five bee families found in Chile based on the dataset obtained from Pontificia Universidad Católica de Valparaíso (PUCV) via GBIF. (E) Cumulative number of species recorded over time for each of the five bee families found in Chile based on the dataset obtained from AMNH. (F) Cumulative number of species recorded over time for each of the five bee families found in Chile based on the dataset obtained from PUCV via GBIF. (G) Total number of unique bee records collected and digitized per year across the full dataset. (H) Total number of wild bee species collected per year across the full dataset.

There were significant differences between the two main sources of data (Figure 1). The database of AMNH and the database from Pontificia Universidad Católica de Valparaíso (PUCV) both had over 3000 unique bee records in the unique dataset (5,941 and 3,560, respectively), but differed considerably in species richness, 243 and 150. There were 135 species found in AMNH database but not PUCV and 42 species found in the PUCV database but not AMNH. The AMNH database had 28 unique species across the whole dataset and the PUCV only five. The main differences were found for Apidae with 71 species in the AMNH database and 37 in the PUCV database, and for Megachilidae with 52 and 21 species, respectively. The two datasets also show considerable temporal differences. In the AMNH database 98% of records were recorded after 1949 and 11% after 1999, the 1980s being the decade with the most unique bee records recorded (1,625). In contrast, the PUCV database had 99% of records post-1949 and less than 1% of the unique bee records recorded after 1999; the decade with the most unique bee records recorded was the 1960s reflecting work by H. Toro and colleagues (1,263). Of the digitized records uneven sampling over time was clearer in the PUCV database than AMNH, particularly between 1950 and 1970 (R^2^ = 0.60 and 0.67). The most even sampling for both datasets was between 1970 and 2000 (R^2^ = 0.86 and 0.88). Regionally, the largest number of unique bee records in the AMNH database come from the Región Metropolitana de Santiago (1,430) and in the PUCV database from Valparaíso (1,479). Highest richness was found in Coquimbo (128) for the AMNH database and Valparaíso (80) for PUCV. The type locality and collection event dataset (Ascher, unpublished), which includes all species, provided additional unique data not found in the other datasets, such as spatial coordinates for 118 species represented among digitized non-type specimen records.

Based on the full species list and regional corrections we report that of the 488 described species known from Chile (Ascher and Pickering 2022; Herrera, unpublished; Montalva, & Ruz), 310 are confirmed as endemic to Chile alone. Furthermore, 86 are endemic to Chile and Argentina, 15 to Chile and Peru, 1 to Chile and Bolivia, 1 to Chile, Argentina, and Peru, 7 to Chile, Peru, and Bolivia, and 3 to Chile, Argentina and Bolivia. Therefore, approximately 423 species (87%) found in Chile are endemic either to Chile alone or shared with neighbouring countries (Ascher and Pickering 2022). When comparing the different bee families present in Chile, the highest single country endemism is found in Andrenidae (92%) followed by Colletidae (80%), Halictidae (56%), Apidae (42%) and Megachilidae (37%). There are 47 species unique to a specific administrative region within Chile, 19 Colletidae, 17 Andrenidae, five Megachilidae, four Halictidae and two Apidae. Regional endemism is focused mainly in two areas of Chile, 24 of the 47 regional endemics are found in the four most northern regions and 11 species are endemic to Coquimbo.

Selection for well-sampled assemblages (used in the GDM model) in Central Chile resulted in 137 5×5 km grid cells from a total of 145 well-sampled cells across the whole of Chile. There was a total of 6,835 unique (date and location) bee records included in these cells from 256 species. Across the five bee families in Chile the unique (per grid cell/site) number of occurrences were distributed as follows: there were 984 grid occurrences of 62 species of Apidae, Colletidae (811, 86), Megachilidae (649, 40), Andrenidae (362, 46), and Halictidae (259, 22). Fifty-seven species were only found in a single well-sampled grid cell.

### GDM

#### Compositional dissimilarity

The strongest abiotic predictors included in the final model, based on the sum of their 3 I-spline functions ((∑i-spline) (the extent to which β-diversity changes along a gradient), was geographic distance (1.57), followed by bare/sparse vegetation cover (0.52), mean diurnal range (mean of monthly (max temp - min temp)) (0.45) precipitation, and annual precipitation (0.26) (Figure 2a). The shape of the splines, and therefore compositional turnover, fluctuated along each of the chosen predictor variables in both magnitude and rate (Figure 2). The final compositional model for Central Chile with abiotic predictor variables and geography explained 34.1% of deviance in compositional dissimilarity and had a mean absolute error of 0.09 ±0.01 (differences between the predicted dissimilarity and the actual dissimilarity). From this deviance, 3.1% was explained by variation along a latitudinal and longitudinal gradient alone, 9.7% by the environmental factors, and the remaining 21.3% was explained by the inclusion of both the spatial and environmental components. The majority of this 21.3% was explained by the interaction between climate and spatial patterns (12.1%) and the combination of all landscape, climate and spatial factors (8.1%). Of the deviance explained by the environmental factors, 45% was attributed to climate, 40% was attributed to climate and 15% by the combination of landscape and climate.

**Figure 2.**
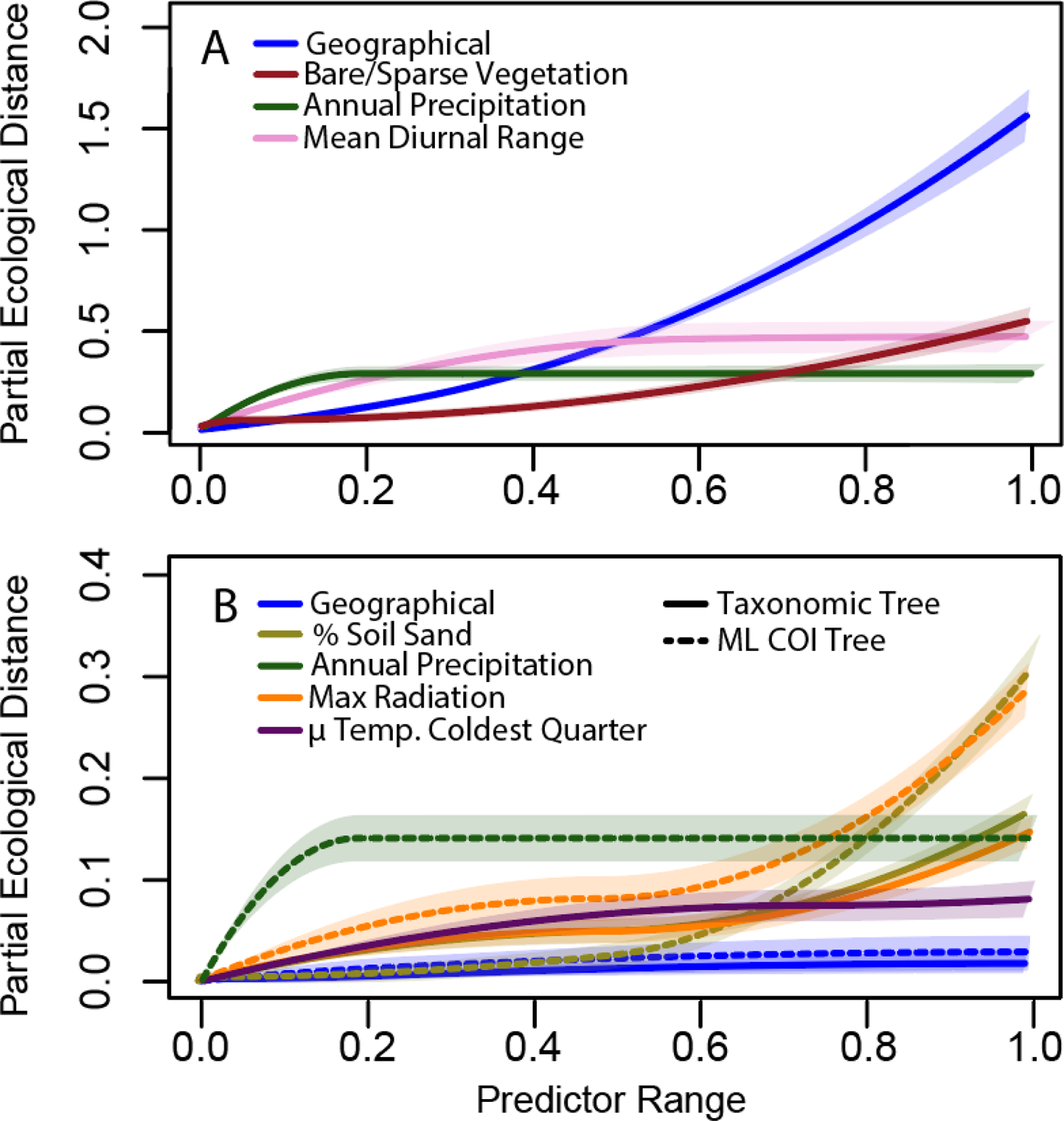
GDM-fitted I-splines (partial regression fits) for variables significantly associated to (A) total β diversity and (B) total phylogenetic diversity. Partial ecological distance refers to the total amount of ecological turnover of β diversity and wild bee phylogenetic diversity. The height of each spline shows how much β diversity or phylogenetic diversity turnover is associated to that variable. Predictor range shows the standardized total range of each gradient across the study area. The predictors shown are those that remained significant at alpha of 0.9 in the backwards elimination models selections. Geographical ranges from −40.7 degrees in the south to −25.3 degrees in the north and −67.2 in the east to −74.1 in the west. Bare/sparse vegetation ranges from 0 to 100% coverage within a 5 x 5 km grid. Annual precipitation (Bio12) ranges from 8mm to 2536mm. Man diurnal range, includes 7.3°C to 16.8°C. Percentage of soil sand ranges from 28.6% to 79.2%. Maximum solar radiation ranges from 19309 kJ m^-2^ day^-1^ to 31334 kJ m^-2^ day^-1^. Mean temperature of the coldest quarter ranges from −13.4°C to 16.2°C. All variables were scaled in the modelling process.

The greatest compositional dissimilarity was found at high values of geographic distance suggesting that well-sampled assemblages in the far south and east varied most from well-sampled assemblage in the far north and west (Figure 2a; Figure 3a & b; Figure S3). The precipitation annual precipitation demonstrated a sharp rise in compositional dissimilarity at low values and then remained stable and equal as precipitation increased, suggesting the major compositional dissimilarity is between assemblages with almost no precipitation (desertic areas north of the central zone) versus assemblages where there is precipitation (Figure 2a; Figure 3c; Figure S3). Mean diurnal range differed most between areas with very low daily variation, in the south and coastal north (northwest), compared to the rest of Central Chile (Figure 2a; Figure 3d; Figure S3). Proportion of bare/sparse vegetation shows considerable gradients between high mountain areas and northern areas and low southern areas and showed a steady increase in compositional dissimilarity as the percentage of bare/sparse vegetation increased (Figure 2a; Figure 3e). For full details of the compositional model see Table S1.

**Figure 3.**
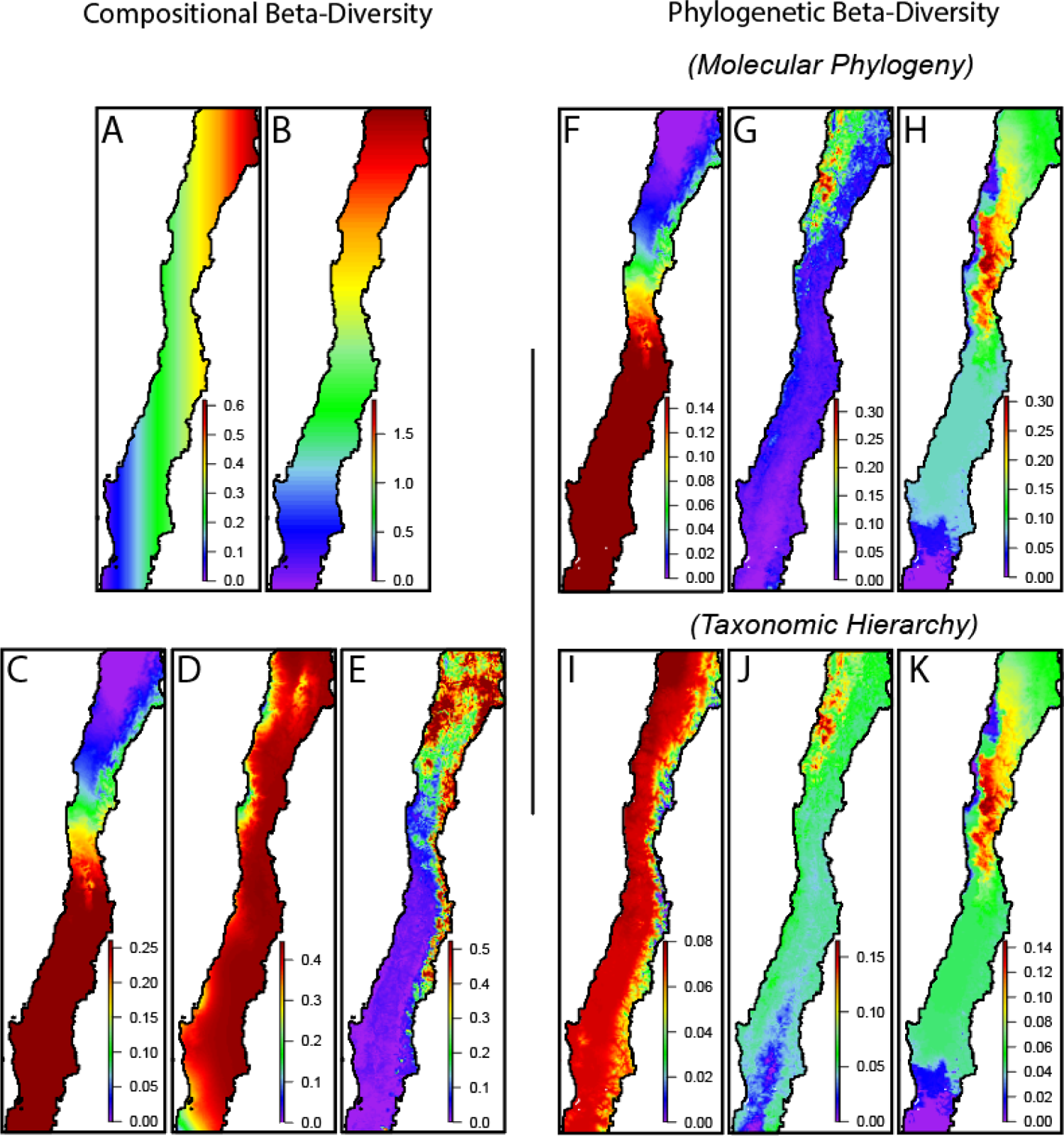
Predictor variables chosen as significant at alpha of 0.9 in the backwards elimination generalized dissimilarity model (GDM) for compositional β-diversity (A-F) and phylogenetic β-diversity (G-K). Colour range shows differences in partial ecological distance, *i.e.*, magnitude of change in β diversity. (A-F) Compositional β-diversity. (A) west to east, (B) south to north, (C) annual precipitation (mm)(Bio12), (D) mean diurnal range (mean of monthly (max temp - min temp)) (Bio2) (°C) (E) Percentage cover of bare/sparse vegetation,, (F-H) Phylogenetic β-diversity calculated using a molecular tree from all publicly available sequences of the mitochondrial gene cytochrome oxidase I (COI). (F) annual precipitation (mm) (Bio12), (F) percentage soil sand content, (H) maximum monthly solar radiation (kJ m^-2^ day^-1)^). (I-K) Phylogenetic β-diversity calculated using a phylogeny based on Linnean taxonomic hierarchy, (I) mean temperature of coldest quarter (°C) (Bio11), (J) percentage soil sand content, (K) maximum monthly solar radiation (kJ m^-2^ day^-1)^). Maps are projected in coordinate system EPSG:31978, SIRGAS 2000 / UTM zone 18S. Not shown is south to north and west to east variables for phylogenetic diversity as they explained very little deviance, see results section for details.

#### Phylogenetic dissimilarity

Overall, environmental variables explained far less deviance of phylogenetic β-diversity than compositional β-diversity for both molecular and taxonomic derived phylogenetic diversity (pβ-diversity). For molecular pβ-diversity, the best model explained only 20.1% of deviance and included soil sand percentage, annual precipitation and maximum solar radiation. Geography alone accounted for only 0.03%, the environmental factors for 13.8 and the remaining 6.2% was explained by the shared combination of spatial and environmental components. Of the limited pβ-diversity explained by the model we observed that increasing soil sand content particularly in the north-west alongside increasing solar radiation from south to north were the key drivers in changes to pβ-diversity (Figure 2b; Figure 3 g, h; Figure S3). Otherwise, low precipitation in the north of Chile (Figure 2b; Figure 3g) explained the observed differences in pβ-diversity (Figure 3f; Figure S3). Even though there was considerable disparity in the number of species in the molecular phylogeny (134) compared to the taxonomic phylogeny (256), the results were broadly comparable. The main difference being the inclusion of mean temperature of the coldest quarter instead of annual precipitation as one of the key factors, which illustrates a distinction in phylogenetic diversity of the high elevation habitats of the Andes (Figure 3i). The taxonomic PD model explained 14.0% of deviance, 0.1% explained by geography alone, 10.7% by environmental factors and 3.2% by the interaction) (Figure 2b; Figure 3i, j, k).

### Assemblage patterns

The k-means clustering of increasing compositional dissimilarity along environmental gradients proposed six ‘bee ecoregions’ (Figure 4). The sites within the WBE strongly reflect geographical patterns. Zona Sur (ZS) reflects the area with the lowest elevation, highest precipitation, lowest temperatures and a low percentage of bare ground. Central Sur (CS) is a continuum from ZS. The Central Andes Region (AC) is characterized by extremely high elevations, lower temperatures, lower solar radiation and higher precipitation than more northern regions and a high percentage of bare/sparse vegetation. The Central Region (ZC) represents an area of more variable conditions, with areas of high elevation, regular precipitation and few areas of bare/sparse vegetation. The Coquimbo Norte Region (CQ) has some areas of medium elevation, low precipitation, high but highly variable solar radiation and an intermediate percentage of bare/sparse vegetation cover. The eastern part of the Atacama Region (AE) is characterised by high to low variation in elevation, very low precipitation, colder temperatures during the coldest period and a high percentage of bare/sparse vegetation.

**Figure 4.**
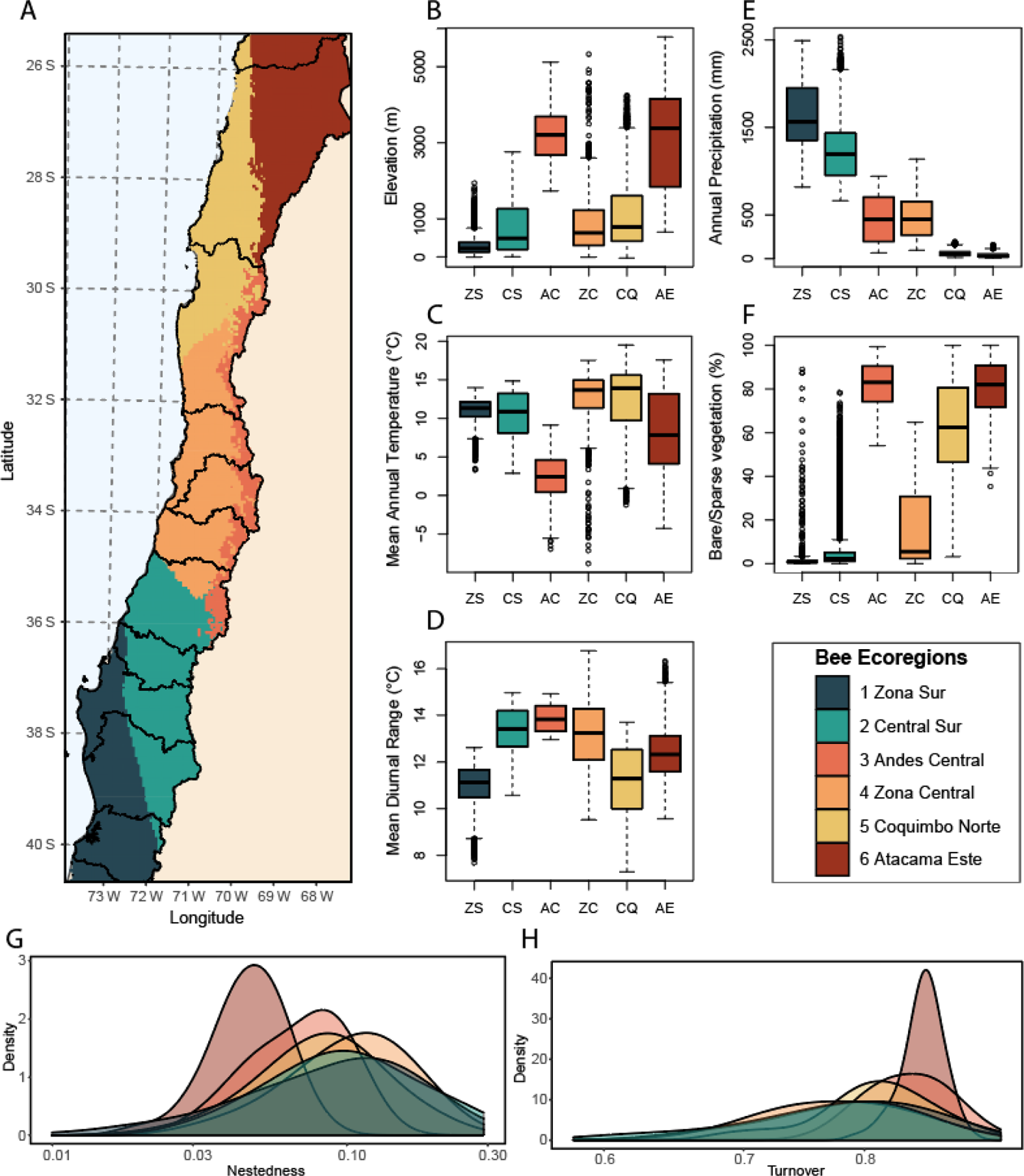
Biogeographical clusters of β diversity turnover along geographic and environmental gradients. (A) The wild bee ecoregions (WBE) (biogeographical clusters) chosen by k-means clustering based on Euclidean distances using multiple indices and taking the mean number of clusters (minus the min and max clusters). The predictors shown (C-F) are those that remained significant at alpha of 0.9 in the backwards elimination models selections. B-F show boxplots of environmental conditions in each of the six WBE. (B) Elevation (m). (C) mean annual temperature (°C; Bio1) (D) Mean diurnal range (mean of monthly (max temp - min temp)) (°C; Bio2). (E) Annual precipitation (mm; Bio12). (F) Percentage cover of bare/sparse vegetation. Overall components of beta diversity, (G) nestedness and (H) turnover, based on 100 random re-samples of 10 pairwise comparisons between sites across all sites with at least three species for each WBE. Map is projected in coordinate system EPSG:31978, SIRGAS 2000 / UTM zone 18S.

Zona Sur, CS and ZC WBE had significantly lower β-diversity turnover across all sites compared to the CQ, AE and CA (all p<0.001). Other notable differences were that the AE and CA regions had significantly higher β-diversity turnover than all other regions (p<0.001) (Figure 4h). Nestedness showed similar patterns, region AE had significantly lower nestedness than all other regions (p<0.0001) (Figure 4g). At the regional and site scales there were noticeable differences in phylogenetic biodiversity, particularly at the limits of our central Chilean study area, and this depending on whether the metric was measured using the molecular phylogeny or the taxonomic hierarchy. The ZC WBE had the highest PSR and PD across both scales and both phylogenetic tree methods, due in part to it having the highest species richness of all bee ecoregions (Table 1). Alternatively, despite its lower species richness the AE bee ecoregion had the highest measures of PSC at the regional scales, indicating that the species present were the most phylogenetically distinct and unique across the WBE (Table 1). Bee ecoregion CQ had high values for all metrics suggesting a rich and phylogenetically diverse WBE with a broad range of Chile’s total phylogenetic diversity.

**Table 1.** Phylogenetic biodiversity measures for each of the wild bee ecoregions (WBE).

| Bee<br>Ecoregion | Molecular Phylogeny |  |  |  |  |  |  |  |  | Taxonomic Hierarchy |  |  |  |  |  |  |  |  |
| --- | --- | --- | --- | --- | --- | --- | --- | --- | --- | --- | --- | --- | --- | --- | --- | --- | --- | --- |
|  | Regional Scale |  |  |  |  | Site Scale |  |  |  | Regional Scale |  |  |  |  | Site Scale |  |  |  |
|  | SR | PSV | PSC | PSR | PD | PSV | PSC | PSR | PD | SR | PSV | PSC | PSR | PD | PSV | PSC | PSR | PD |
| 1. Zona Sur (ZS) | 55 | 0.85±0.004 | 0.34 | 46.93±0.21 | 13.80 | 0.83±0.15 | 0.67±0.21 | 5.08±4.96 | 2.00±1.69 | 110 | 0.71±0.01 | 0.035 | 78.14±1.15 | 10.46 | 0.66±0.16 | 0.27±0.19 | 5.93±6.71 | 3.18±1.44 |
| 2. Central Sur (CS) | 66 | 0.86±0.003 | 0.32 | 57.08±0.19 | 14.84 | 0.83±0.11 | 0.63±0.15 | 6.54±5.79 | 2.55±1.90 | 129 | 0.74±0.01 | 0.035 | 94.63±1.19 | 11.34 | 0.67±0.15 | 0.23±0.18 | 7.90±7.95 | 3.82±1.51 |
| 3. Andes Central (AC) | 60 | 0.87±0.003 | 0.35 | 51.98±0.20 | 14.36 | 0.78±0.21 | 0.59±0.20 | 6.88±5.80 | 2.72±2.04 | 109 | 0.73±0.01 | 0.036 | 79.12±1.15 | 10.41 | 0.62±0.12 | 0.18±0.11 | 8.24±7.39 | 3.71±1.59 |
| 4. Zona Central (ZC) | 100 | 0.86±0.001 | 0.28 | 86.43±0.14 | 21.53 | 0.83±0.09 | 0.62±0.15 | 7.71±6.49 | 2.99±2.28 | 211 | 0.73±0.01 | 0.026 | 154.06±1.27 | 14.12 | 0.66±0.14 | 0.21±0.16 | 8.60±7.99 | 3.92±1.74 |
| 5. Coquimbo Norte (CQ) | 87 | 0.87±0.002 | 0.30 | 76.04±0.16 | 19.73 | 0.84±0.10 | 0.68±0.15 | 6.49±6.09 | 2.60±2.14 | 162 | 0.73±0.01 | 0.032 | 118.20±1.25 | 12.96 | 0.70±0.15 | 0.27±0.21 | 7.63±7.19 | 3.75±1.63 |
| 6. Atacama Este (AE) | 33 | 0.87±0.007 | 0.39 | 28.54±0.23 | 9.18 | 0.71±0.22 | 0.61±0.21 | 3.07±2.53 | 1.31±1.03 | 63 | 0.71±0.02 | 0.046 | 45.68 ±0.98 | 8.46 | 0.64±0.16 | 0.30±0.14 | 4.18±2.89 | 2.85±1.09 |

Molecular Phylogeny refers to the tree based on a molecular phylogeny, which was produced based on all publicly available sequences of the mitochondrial gene cytochrome oxidase I (COI) available on GenBank and has fewer species but more accurate measures of branch lengths. Taxonomic Hierarchy refers to the tree based on the Linnean classification using Rank, Family, Subfamily, Tribe, Subtribe, Genus, Subgenus, Species and includes all species but less accurately measures branch lengths. SR= Species richness, total number of species included in the tree for that ecoregion. PSV = Phylogenetic species variability; the extent of relatedness between species in a community. PSC = Phylogenetic species clustering; a measure of the distance between branch tips across a community phylogeny. PSR = Phylogenetic species richness; PSV multiplied by the species richness. PD = Phylogenetic Diversity; the sum of the total branch length. At the regional scale the plus-minus sign (±) indicates the expected standard deviations per community, at the site scale it is the standard deviation of the values across sites.

Of the 336 (79% of all georeferenced bees we had for Chile and approximately 69% of all known Chilean bees) bee species recorded in Central Chile from our final database of all unique bee records, were endemic to Chile only. Each WBE had a high degree of endemism (Table S2) and was defined by the presence of some key indicator species which was disproportionately likely to be found across multiple sites within specific WBE (Table S3). The highest species richness was recorded from ZC with 211 species, of which 137 (68%) were endemic to the country; and 39 of these endemic species were unique to this ecoregion. Four species were calculated to be indicative of ZC including *Anthidium chilense* Spinola, 1851 (Hymenoptera: Megachilidae) (Table S3). The second most species rich area was CQ with 162 species recorded and 113 endemic species (73%), the highest of all WBE, and of which 25 were unique to CQ. Seventeen species were indicative of the CQ region, the most statistically significant were *Neofidelia profuga* Moure & Michener, 1955 (Hymenoptera: Megachilidae) and *Chilicolletes delahozii* (Toro, 1973) (Hymenoptera: Colletidae). Region CS had a total of 129 species recorded, 63 were endemic (52%) and of those endemic, 21 unique. *Corynura patagonica* (Cockerell, 1919) (Hymenoptera: Halictidae) was the most statistically significant of the six species indicative of the CS region. Region ZS had 110 species and the lowest endemism with 55 species (51%,), 12 of which were unique. The region was indicated by the presence of *Diphaglossa gayi* Spinola, 1851 (Hymenoptera: Colletidae) *Corynura corinogaster* (Spinola, 1851) (Hymenoptera: Halictidae). Region AC had an observed species richness of 109 species, with 58 endemics (57%), and of which 10 were unique. *Alloscirtetica gazullai* (Ruiz, 1938) (Hymenoptera: Apidae)*, Anthidium chubuti* Cockerell, 1910 (Hymenoptera: Megachilidae), and *Anthidium espinosai* Ruiz, 1938 (Hymenoptera: Megachilidae) were the most indicative out of 15 species. Finally, region AE had the lowest recorded richness of 63 species, 35 endemics (67%, five unique); 3 species were indicative of the region including *Callonychium atacamense* Toro and Herrera, 1980 (Hymenoptera: Andrenidae), and *Chilicola travesia* Toro and Moldenke, 1979 (Hymenoptera: Colletidae), (see Figure 5 and S4 for full details of species diversity in each WBE).

**Figure 5.**
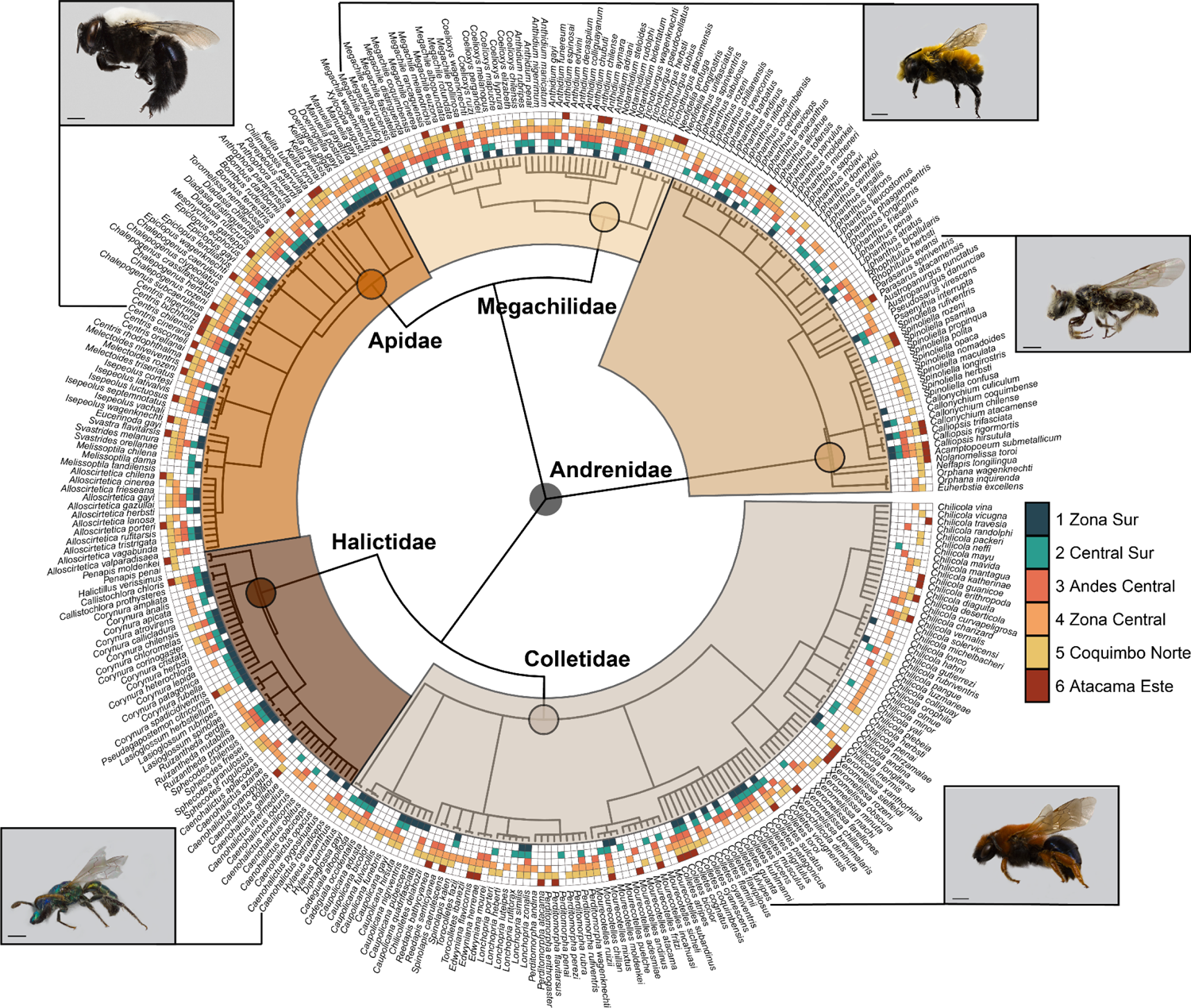
Tree (taxonomic hierarchy) of all unique bee records within Central Chile. Coloured squares show the ecoregions (Figure 4) where a species was historically collected. Illustrated are single species from each family of wild bees found in Central Chile, from top right going clockwise are illustrated *Megachile semirufa* Sichel, 1867 (Hymenoptera: Megachilidae)*, Rhophitulus evansi* (Ruz & Chiappa, 2004) (Hymenoptera: Andrenidae). *Colletes fulvipes* Spinola, 1851 (Hymenoptera: Colletidae)*, Caenohalictus rostratriceps* (Friese, 1916) (Hymenoptera: Halictidae), and *Centris cineraria* Smith, 1854 (Hymenoptera: Apidae). All photographs by Patricia Henríquez-Piskulich, Instituto de Entomología, Universidad Metropolitana de Ciencias de la Educación, Santiago, Chile.

## Discussion

### What are the main drivers of wild bee taxonomic and phylogenetic diversity in Chile?

Here we describe the large-scale distribution patterns of wild bees in Chile and their underlying drivers. Whereas β-diversity of wild bee assemblages in Chile represents high turnover across the country, reflecting its unique and varied geography, by contrast Phylogenetic β-diversity showed overall greater homogeneity across the country, suggesting that while assemblages changed in composition, they generally encompassed similar levels of phylogenetic diversity. Generalized dissimilarity modelling showed that along latitudinal and longitudinal gradients, the make-up of wild bee assemblages became increasingly different from each other, particularly from south to north. This turnover across space, measured as geographic distance, accounts for obvious distinctions in habitat, climate and topography with different landforms, bioclimatic zones and ecoregions (Dinerstein et al., 2017, Sarricolea et al., 2017). Indeed S-N latitudinal variation was correlated to increases in temperature and decreases in precipitation and changes in ecosystems. The study area chosen for this research included many different habitat types and bioclimatic zones, ranging from temperate to semi-arid and sub-tropical habitats and Mediterranean like climates to high altitude Tundra like climates (UGS 2010, Sarricolea et al., 2017).

Following latitudinal patterns, west (Pacific coast) to east (Andes) changes in elevation and climate drive β-diversity turnover as well. Increases in elevation from the coast to the mountain ecosystems in this case, imply sharp changes in relatively short horizontal distances in the characteristics of the physical environment (Sundqvist et al., 2013). These variations drastically affect the composition of several organisms, including wild bee assemblages (Hoiss et al., 2012; Orr et al., 2021a). In this study, higher elevations corresponded to areas with a lower minimum temperature in the coldest periods, which are likely to favour species with weaker competitive strength but greater adaption to this extreme environment, because of an increased pressure of environmental filtering rather than competition in these high elevation environments (Hodkinson, 2005; Hoiss et al., 2012).

Proportion of bare/sparse vegetation was the only habitat variable chosen in the final model. High percentages of bare/sparse vegetation in Central Chile occur in the high Andes and the arid landscapes of southern Atacama. Bare/sparse vegetation and associated soil conditions offer key nesting resources for certain species of wild bees (Cane 1991; Danforth et al., 2019). High proportions of bare soil are found across a wide elevation and latitudinal gradient, and therefore interact with temperature and vegetation in terms of wild bee habitat. Bare/sparse vegetation correlates to other predictors, including floral diversity and composition, and soil characteristics which are likely to be important drivers of wild bee diversity at the fine scale (Michener 2000, Murray et al., 2009, Rodríguez et al., 2021). Precipitation was less important overall but vital in distinguishing wild bee assemblages found in desert regions. The unique and lower bee diversity observed in desert areas in Chile is likely due to localized and unpredictable precipitation in these areas, which only allows for the persistence of species capable of multi-year diapause and thus well adapted to these conditions (Moldenke 1976b; Danforth et al., 2019).

The ability of our model to capture the β-diversity patterns of Chilean wild bee assemblages measured as the percentage of explained deviance in our compositional model is 34%, which is lower than some other studies on insect groups (Chesters et al., 2019), but it is in line with results from areas with considerable turnover and limited well-sampled sites (Laidlaw et al., 2016). Stochastic, historical and evolutionary processes, which were not captured by our models, might also drive compositional dissimilarity (Wiens and Donoghue 2004, Ricklefs 2008, Warren et al., 2014). It is difficult to draw any meaningful conclusions on the drivers of pβ-diversity. At the spatial resolution of this study, geographic, habitat and climate variables explain only a moderate amount of the variation in phylogenetic β-diversity. However, we observed that the most important variables reflected those of compositional β-diversity and that there is variability in some measure of phylogenetic biodiversity within the proposed WBE. Particularly the north and the north-east of the study region show high turnover in pβ-diversity. The well-sampled assemblages used to train the GDM were only a subset of the full diversity found in Central Chile and it is possible this selection impacts the GDM and resulting WBE. However, a broad range of the diversity and endemism of Central Chile was included in the well-sampled assemblages. For example, lineages which are endemic to Chile at the tribal rank, Penapini and Nolanomelissini, are included in the GDM analysis as part of well-sampled assemblages. The two *Penapis* Michener, 1965 species included within well-sampled assemblages, *Penapis moldenkei* Bohart, Toro, and Rozen, 1997 (Hymenoptera: Halictidae) (AE + CQ) and *Penapis penai* Michener, 1965 (Hymenoptera: Halictidae) (CQ), are both indicator species for the two northern WBE. *Nolanomelissa toroi* Rozen, 2003 (Hymenoptera: Andrenidae) is also representative of the CQ WBE. The inclusion of these highly distant taxa will have improved the GDM representation of the totality of Chilean wild bee diversity and their role as indicator species gives support to the conservation relevance of the WBE described here.

### Can we classify *de novo* ecoregions of wild bee diversity (WBE) using their occurrences and environmental conditions and are these WBE consistent with biogeographical regionalisations based on other organisms?

The WBE presented here are an important addition to understanding biogeographic patterns in Chile. We present for the first time a statistical representation of regional variation in wild bee assemblages, reinforcing patterns already observed in the field for Chilean bees (Moldenke 1976b) and both supporting and contrasting biogeographic distinctions made for other taxa (Morrone, 2015). The WBE reflect the main climate and habitat drivers outlined above, with similarity to the bioclimatic zones of Chile (Sarricolea et al., 2017). Furthermore, the most northern and southern WBE show the lowest diversity, giving support to the Central Chile distinction made when selecting the study area. The transition zones between WBE as proposed here show similarity with those proposed by Morrone (2006 & 2015) but more closely reflect the finer scale districts proposed by Peña (1966) based on the diversity of Tenebrionidae (Coleoptera).

The transition between the ZS WBE and the CS WBE reflects, at the broadscale, the transition from the Subantarctic subregion to Central Chilean subregion (Morrone, 2015). The colder and more humid ZS WBE is also closely aligned to the Valdivian Forest Region as categorised by Kuschel (1960) and Peña (1966). The rest of WBE are found within the Central Chilean subregion and South American transition zone (Morrone, 2015). Within the central Chilean Subregion there are 3 WBE which differ from previously described biogeographic regions. It is a global hotspot for the diversity of *Colletes* Latreille, 1802 species (Bystriakova et al., 2018), with many species likely endemic to the subregion (Ferrari, 2017). The CS ecoregion overlaps considerably with Central Valley (Morrone, 2015; Peña, 1966) which it shares with ZC, but unlike previous regionalisations, the wild bee assemblages do not show a distinction at the Southern Andean Cordillera or the coastal desert (Peña, 1966), instead presenting a single assemblage across all three districts. The transition to ZC also reflects northern limit of conifer plantations (Martínez-Tilleria et al., 2017).

The ZC along with the CG WBE is characteristic of the Mediterranean biodiversity hotspot in Chile and the high diversity observed reflects global diversity patterns of wild bees, such as a preference for non-forest habitat, reported by Linsley (1958) and Michener (1979), and recently modelled by Orr et al., (2021a). The ZC includes a transition zone between ecosystems which separates the other two WBE (CS and CQ) within the Central Chilean subregion from Lower Montane Forest to Mediterranean Xeric Shrub (USGS, 2010). The 137 endemic species found in this WBE support the hypothesis that Mediterranean regions have been a centre of speciation (Moldenke 1976b, Danforth et al., 2013). The CQ WBE reflects the transition between biogeographic provinces, Santiago and Coquimbo (Morrone et al., 2006), which corresponded to the transition zone between xeric and sclerophyllous biomes (USGS, 2010). It includes the intermediate desert district and the Coquimban desert as well as following the northern coast district northwards(Peña, 1966) and strongly reflects the evergreen scrub with microphyllous leaves, terrestrial ecosystem proposed by Martínez-Tilleria et al., (2017). The CQ WBE represents high phylogenetic diversity overall and has the highest number of indicator species including endemic tribal lineages (*Penapis penai and Nolanomelissa toroi*), as well as highly distinct taxa such as *Orphana wagenknechti* Rozen, 1971 (Hymenoptera: Andrenidae), and *Neofidelia profuga* both of which are endemic to the CQ WBE. These three WBE in the Central Chilean Subregion represent a favourable climate for bee species However, the geographical isolation of the subregion has yielded a lower species richness than in similar environments such as in California which is home to over 1,600 species of bees (Moldenke 1976a, Moldenke 1979, Hung et al., 2019) and the Mediterranean (Nieto et al., 2014; Leclerqc et al., 2022).

Towards the north of Central Chile, CQ transitions to the AE WBE which is consistent with the transition zones of the Intermediate Desert to the Northern Desert as well as from the Central Andean Cordillera to Northern Andean (Peña, 1966: Morrone, 2015). The CQ and AE WBE to the north are defined by very low precipitation but differ in elevation and temperature, and therefore in the identity and timing of flowering plants. The unique endemics from these regions include several species from the diverse and endemic genus *Chilicola* Toro & Moldenke, 1979 (Hymenoptera, Colletidae), whose divergences, determined through molecular dating, reflect the latitudinal moisture gradient, as well as the previous biogeographic zone’s descriptions (Monckton 2016; Morrone, 2015), supporting the distinctions made by the GDM. The AE WBE also shows high values of PSC suggesting it is the region with greatest evolutionary diversity, even though it has a lower species richness.

The east-west transition to the Central Andes bee eco-region follows the transition zone between the Central Chilean Subregion and the South American transition zone, and follows the Central Andean Cordillera proposed by Peña (1966), with a highly distinctive fauna and vegetation (Arroyo et al., 1982). The Chilean Andes are, almost certainly, home to many undescribed and undigitized bee species especially given the high diversity found in other parts of the Andes (Henríquez-Piskulich et al., 2020, Gonzalez and Engel 2004). This is supported by two recently described *Colletes* species endemic to the Central Andes WBE; *Colletes flavipilosus* Ferrari, 2017 (Hymenoptera: Colletidae), and *Colletes guanta* (Ferrari 2017). This limited knowledge of the diversity of unique endemic species is also similar for the far north (AE), where the availability of digitized records regarding the diversity of wild bee species is much lower than for the Mediterranean Central Zone.

There is still considerable turnover, unique diversity and localized endemism within all the proposed WBE, and this is likely because of conserved traits among species that facilitate their persistence in varying habitats and climates (Villalobos and Vamosi 2018) but may also reflect a lack of sampling and digitization in the understudied areas. There is preliminary evidence for a latitudinal and elevation relationship in rates of turnover; the northern and eastern WBE both for Central and Northern (CQ, CA, AE), show higher turnover between sites compared to the more southern, lower elevation WBE, reflecting a higher exchange and filtering of species (Hodkinson, 2005; Hoiss et al., 2012) as has been observed in for bees in the central Chilean Andes (Henríquez-Piskulich et al., 2020). In contrast to the results observed for compositional dissimilarity, we did not observe such a strong variation in phylogenetic diversity metrics between WBE. One explanation for this is that because of Chile’s long isolation, wild bee lineages have been able to occupy a wide variety of habitats across the country and this is reflected in the considerable radiations of genera with species endemic to Chile (Moldenke 1976b), for example, *Liphathus* species and Xeromelissine bees (Monckton, 2016; Packer & Ruz, 2016). There is, however, some variation in phylogenetic biodiversity metrics at the site level, suggesting there is a finer scale gradient of land use, floral diversity and nesting resources that may explain phylogenetic diversity, as has been observed comparing bee phylogenetic diversity in urban, forest and agricultural ecosystems in north-eastern USA (Harrison et al., 2018).

### How do these classifications represent the conservation threats towards Chile’s unique wild bee fauna?

Wild bees in Chile face similar threats as wild bees globally, including large-scale anthropogenic land use conversion and intensification, climate change, as well as the introduction of pathogens and non-native species (Potts et al., 2016; Morales et al., 2013). Nonetheless, these threats vary regionally. The WBE classification presented here highlights the threats unique assemblages are likely to face. The Mediterranean-type region of Chile is densely populated, has a long history of agriculture and only a small percentage of its area is protected (Underwood et al., 2009). Agroecological transitions, involving optimization of management and land use alterations, are probably necessary to ensure coexistence between agriculture production and native wild bees, in particular ensuring that natural scrub habitats are not removed (Henríquez-Piskulich et al., 2021; Hung et al., 2022; Rodríguez et al., 2021). Furthermore, according to Petit et al., (2018) most protected areas in the Central Mediterranean-type biome and the arid desert north of Chile do not have a clear management plan, which threatens biodiversity conservation. High elevation areas such as the Andes also face threats from human-induced land use change, including mining activities, and grazing, and are also likely to experience the impacts of climate change sooner than other areas (Hodkinson 2005, IPCC 2014). For instance, the presence of human-introduced exotic weeds altered the structure of wild bee assemblages in Farellones in the Chilean Andes, particularly affecting large and medium-sized bee species (Henríquez-Piskulich et al., 2018). In addition, climate change is anticipated to cause species to shift to higher elevations (Parmesan and Yohe 2003, Marshall et al., 2020), increase competition between species and increase the chance for temporal mismatches between wild bees and the plant resources they pollinate (Hegland et al., 2009).

Even though we use an unprecedented dataset to show robust biographical patterns of bees in Chile, there are clear shortfalls and gaps in our knowledge which hamper a full understanding of wild bee diversity, distribution and ecology in Chile (Cardoso et al., 2011). The Linnean shortfall (undescribed species) is visible in the difficult groups. Several Halictidae species were excluded from the GDM analysis because of the difficulty in determining species, limiting their digitization. Additionally, there are many collected and sorted morphospecies in the AMNH collection awaiting description, many likely representing distinct and important biodiversity. The Wallacean shortfall (unknown distributions) is visible in the many species whose distribution we can only estimate from a handful of geo-referenced occurrence points, this includes distinct taxa which were classified as regionally endemic and/or indicator species. Finally, the ecology, such as thermal tolerances (Hutchinsonian Shortfall) and biotic interactions (e.g. floral prefence) (Eltonian Shortfall) (Hortal et al., 2015), of many of these species is poorly described or unknown. The filling of these gaps and shortfalls requires; (i) the support and training of experts in Chilean bee taxonomy alongside modern technical approaches (Engel et al., 2021; Orr et al., 2021b) (ii) the funding of local and regional inventories, similar to the cooperative expeditions of ANMH and local Chilean researchers, (iii) the identification, digitization and sharing of specimens in existing collections, and (iv) life-history focussed studies alongside modern techniques such as pollen DNA barcoding (Bell et al., 2016). These data gaps are universal, particularly for vast groups of organisms like insects. Using available data and testing ecological theories is necessary for a better use and understanding of currently available information (Diniz-Filho et al., 2010). An important step in biodiversity conservation is to take monitoring and inventory data and use this to understand regional patterns in diversity (Socolar et al., 2016), which is what we have done here. The *de novo* WBE presented here not only improves our understanding of biogeographic patterns but has clear potential in supporting conservation efforts.

Ecoregions can highlight priority areas with distinct or high-value biodiversity areas worthy of greater attention, as well as showing where threats to biodiversity are most extreme (Olson et al., 2001; Watson & Venter et al., 2017), and if protection is evenly distributed (Dinerstein et al., 2017). The indicator species highlighted here for each WBE can be effective examples of distinct and high-value biodiversity. For example, *Penapis penai* and *Nolanomelissa toroi* are tribal endemic species, and are representative of the CQ WBE and can therefore be used to raise awareness of the important desert shrubland and scrubland habitat where they occur, and to draw attention to them as conservation targets. Especially in the case of *N. toroi* which is phylogenetically unique but also highly specialised in its feeding behaviour collecting pollen from *Nolana* species (Rozen, 2003). Equally important are species known to be endemic to the different WBE, such as *Orphana wagenknechti* (CQ) endemic to xeric scrubland habitat or *Eucerinoda gayi* (Spinola, 1851) (Hymenoptera: Apidae) (ZC) endemic to Mediterranean type habitat of Central Chile. Early diverging species such as *Eucerinoda gayi* (Freitas et al., 2022) also represent unique, endemic evolutionary diversity and classifying its habitat, community and distribution at the broader scale should encourage its inclusion in conservation efforts. The GDM and associated ecoregions show where we have environmental correlates to unique and important endemic taxa and phylogenetic diversity versus areas where we still need basic exploration, descriptions and digitization. Here, we show clearly how the transition zone between lower montane forests to xeric scrub and shrubland is essential for the diversity of bees. The ZC and CQ WBE represent a highly endemic and varied fauna in Central Chile and high-quality, natural, habitat unaffected by the conservation threats within Central Chile should be highlighted and conserved. For example, our data show that the natural scrubland south of Vicuña in the Elqui Province, Coquimbo Region, is a clear hotspot of unique bee diversity, highlighting an area that is in unlikely to be included as key area for wildlife conservation but appears to be important for wild bee diversity. Ecoregions defined by turnover in assemblages can be used as proxies for community and species level biodiversity, particularly where high-quality observation data is lacking, and provide a clear outline as to the where conservation should be focussed (Smith et al., 2018). Conserving sites with high species richness and high endemism across WBE in Chile could ensure that additional phylogenetic diversity, and functional diversity, will be conserved (Winter et al., 2013). We propose that our data can be used by the IUCN - Wild Bee Specialist group as a starting point for including Chile and its WBE in efforts to establish threat status and red lists for South American bee fauna. Not only do we present unique assemblages across Chile, but we highlight the taxa representative and endemic to each WBE. Providing a baseline list of species with limited ranges, potentially vulnerable to anthropogenic changes. Analyses of population trends over time may be possible for areas which have been sampled consistently, e.g., in and around Santiago and Valparaiso, (Boyd et al., 2022). Furthermore, we show the relevance and importance of using community science sites such as iNaturalist, which produce an ever-increasing set of validated records and constantly facilitate the detection of new species, for example a recent third *Hylaeus* species adventive in Chile, the recently detected Palearctic species *H. leptocephalus* (Morawitz, 1870) (https://www.inaturalist.org/observations/132205886).

Our results demonstrate how combining data from different institutions, attending to data quality directly by engaging leading identifiers and digitizers, and visualizing patterns and trends corroborate the uniqueness of the Chilean fauna and highlight regions of high endemics and turnover that can be priorities for bee conservation.

## Acknowledgements

LM and NJV were funded by the FNRS/FWO joint programme ‘EOS – Excellence Of Science’ for the project ‘CliPS: Climate change and its impact on Pollination Services (project 30947854). LM was additionally supported by a F.R.S.-FNRS fellowship “Chargé de recherches”. JA was funded by Singapore NRF2017NRF-NSFC001-015. CV was funded by APEX DIUMCE 2019 “Efectos de la Intensificación de la Agricultura sobre el ensamble de Abejas Nativas de Chile y su importancia para los Servicios de Polinización” (project PGI 02-2019). We are grateful to all those involved in collecting, managing, curating, digitising, correcting and sharing bee collection data from Chile that makes this work possible. We would also like to thank Dr. D. Sponsler and Dr. M. Fitzpatrick for their assistance with generalized dissimilarity modelling. We thank Jerome G. Rozen, Jr., and the late Robert G. Goelet who supported bee research at the AMNH, and Laurence Packer of York University shared helpful specimens and unpublished data. Digitization of bees at the AMNH by JA was funded in part by an NSF-BRC grant Collaborative databasing of North American bee collections within a global informatics network project.

## Statement of authorship

NJV, CV, JSA, AB, and LM conceived the study; AB, LM and NJV designed the methodology. JSA, CV, PH, AV, VH and NJV collected, organised and managed the data. LM and AB analysed the data. LM led the writing of the manuscript. All authors were involved in editing the manuscript.

