## Supplementary material for "Chilean bee diversity: Contrasting patterns of species and phylogenetic turnover along a large-scale ecological gradient": Figure S

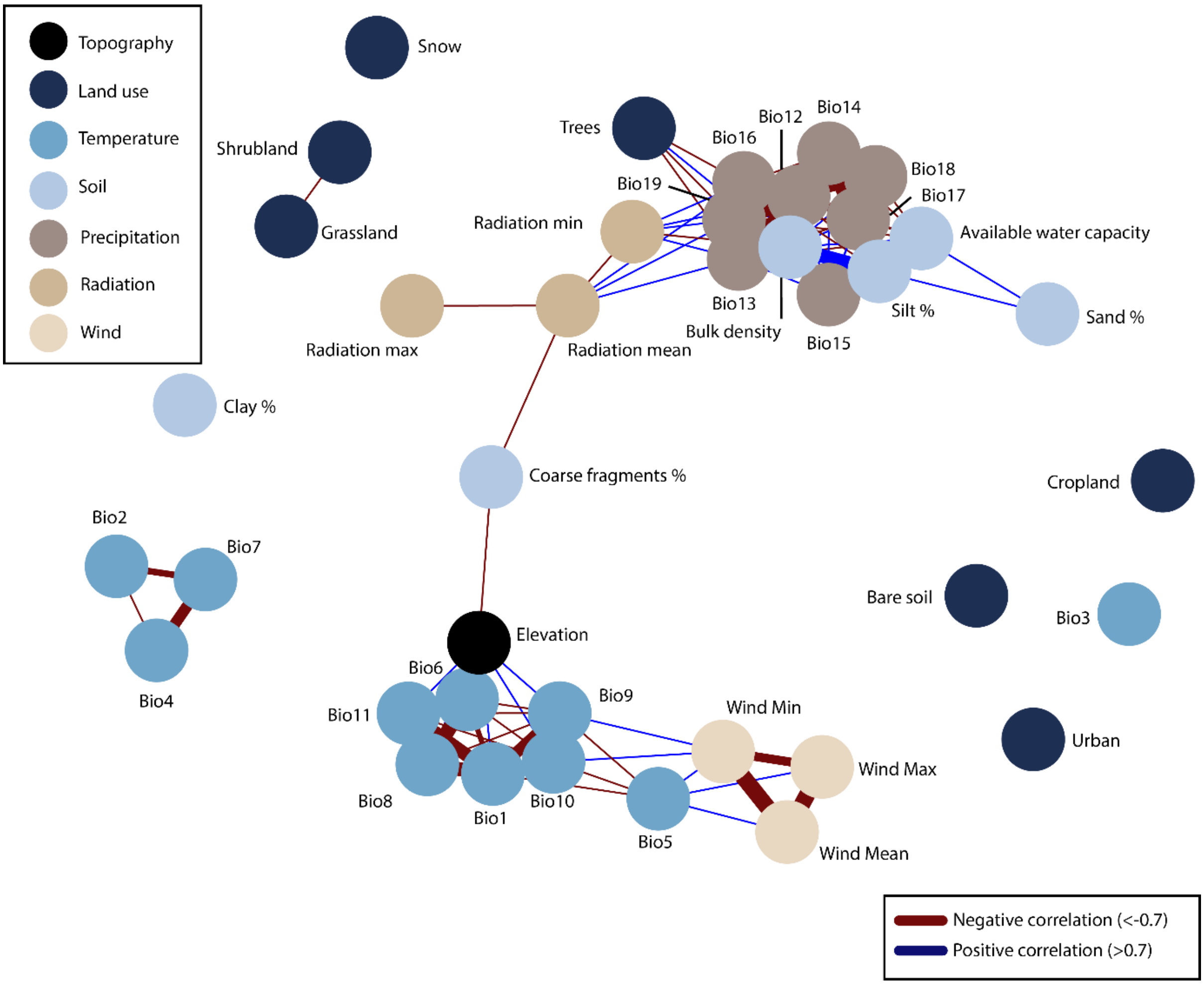


Figure S1. **Correlations (Pearson's correlation coefficient) between all variables at the beginning of the analysis (40).**

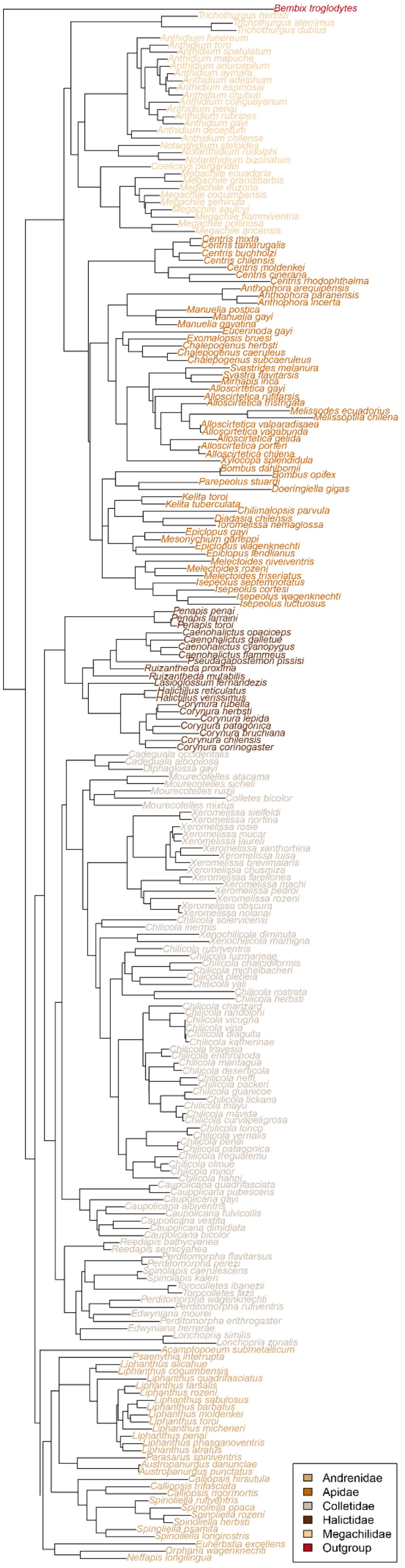


Figure S2. **Maximum likelihood phylogenetic reconstruction based on all publicly available sequences of the mitochondrial gene cytochrome oxidase I (COI) available on GenBank.**

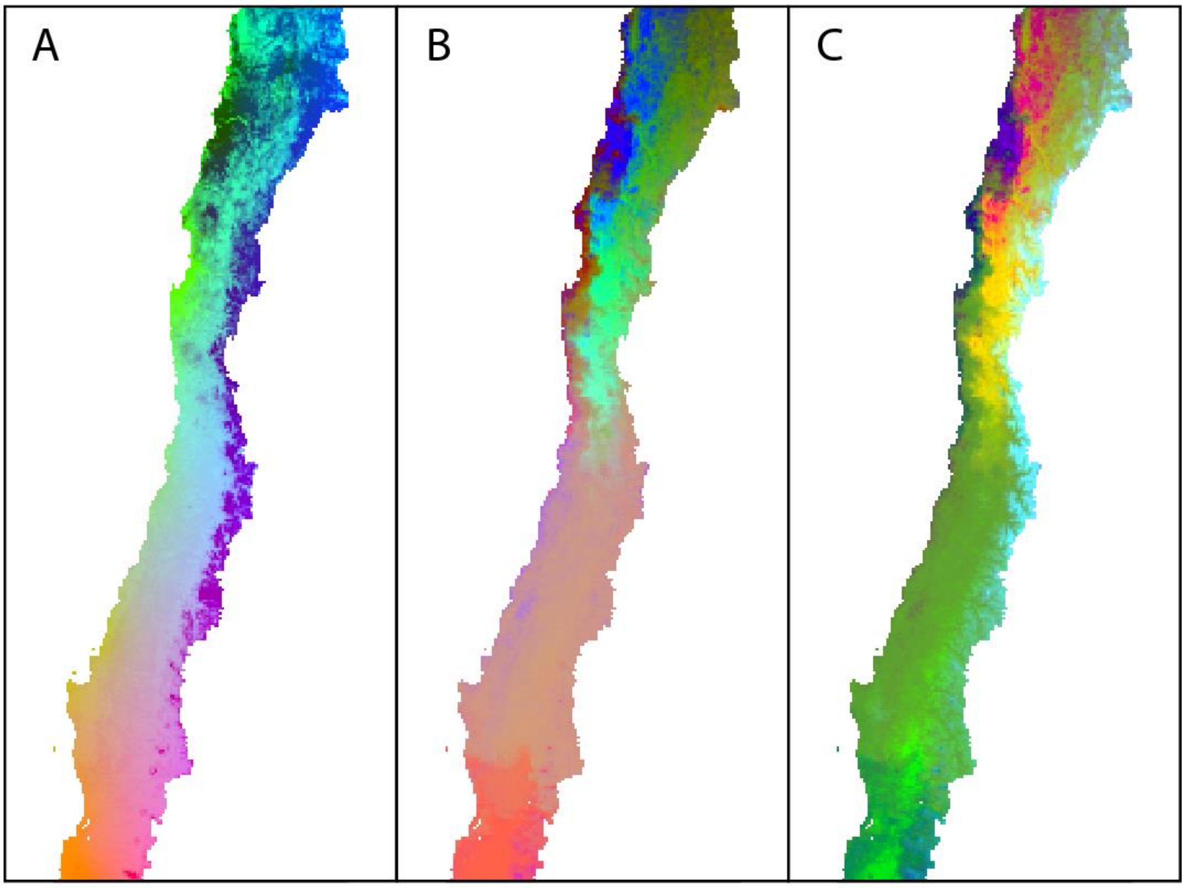


Figure S3. **Predicted spatial variation in wild bee assemblages and phylogenetic diversity.** Colors represent gradients in species composition derived from transformed environmental predictors. Locations with similar colors are expected to have similar assemblages. (A) Species assemblages (compositional diversity). (B) Phylogenetic diversity based on molecular phylogeny of COI. (C) Phylogenetic diversity based on the taxonomic hierarchy of Linnean classification.

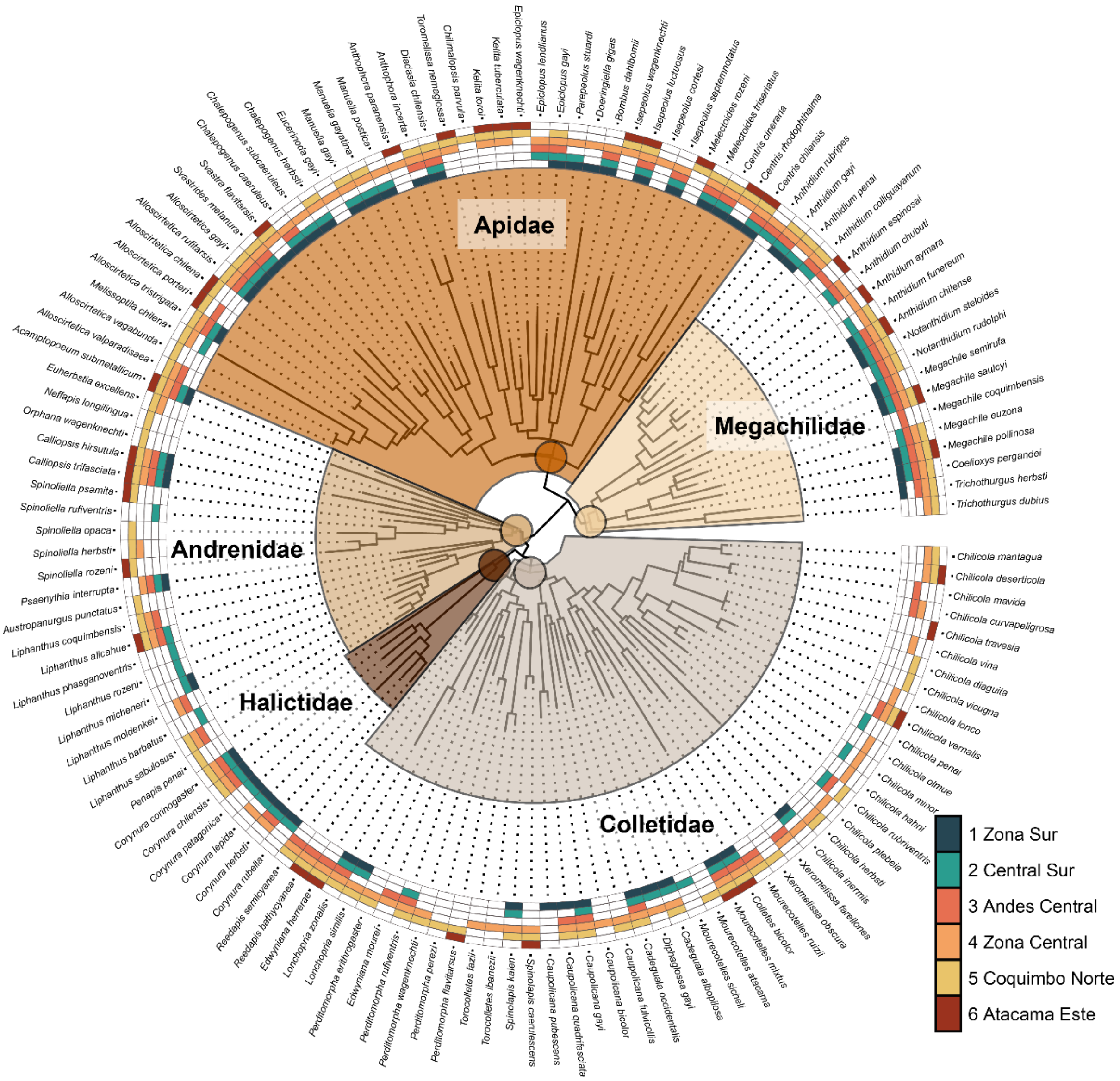


Figure S4. **Tree (molecular phylogeny COI) of all unique bee records within Central Chile for which we had molecular data.** Coloured squares show the ecoregions (Figure 4) where a species was historically collected.

Table S1. **Model output of final compositional GDM used for assemblage predictions.**

| Model Summary | | Model Performance | | |
|---|---|---|---|---|
| NULL Deviance | 308.01 | Mean Absolute Error | 0.09 | ±0.01 |
| GDM Deviance | 203.00 | RMSE | 0.11 | ±0.01 |
| % Deviance Explained | 34.09 | Equalised RMSE | 0.14 | ±0.01 |
| Intercept | 0.94 | Mean Corr. Obs. Pred. | 0.55 | ±0.09 |

**Predictor Summary**

| | *Geographic* | *Bare Vegetation (%)* | *Mean Diurnal Range (°C)* | *Annual Precipitation (mm)* |
|---|---|---|---|---|
| Splines | 3 | 3 | 3 | 3 |
| Coefficient | 0.05 | 0.03 | 0.42 | 0.26 |
| Coefficient[2] | 0.37 | 0 | 0.03 | 0 |
| Coefficient[3] | 1.15 | 0.49 | 0 | 0 |
| Sum of coefficients | 1.57 | 0.52 | 0.45 | 0.26 |

Model performance values are calculated as the average of 1000 cross-validations with 20% of the site data used as test data. RMSE = Root Mean Square Error, Mean Corr. Obs. Pred = Pearson's correlation coefficient for the observed values of dissimilarity against the predicted values. Geographic refers the dissimilarity along latitudinal and longitudinal gradients. Mean diurnal range is measured as the mean of monthly max temp - min temp.

Table S2. **Species endemic to a single wild bee ecoregion (WBE) based on all unique bee occurrences.**

| WBE | Species | #Occurrences |
|---|---|---|
| **1** | *Caenohalictus opacus* | 1 |
| | *Callonychium culiculum* | 1 |
| | *Caupolicana pubescens* | 1 |
| | *Chilicola gutierrezi* | 1 |
| | *Chilicola longitarsa* | 1 |
| | *Chilicola michelbacheri* | 1 |
| | *Corynura ampliata* | 2 |
| | *Corynura heterochlora* | 1 |
| | *Lasioglossum herbstiellum* | 1 |
| | *Lasioglossum rubripes* | 1 |
| | *Liphanthus atratus* | 1 |
| | *Sphecodes friesei* | 1 |
| **2** | *Caenohalictus monilicornis* | 1 |
| | *Chilicola mayu* | 1 |
| | *Chilicola mirzamalae* | 1 |
| | *Chilicola penai* | 1 |
| | *Halictillus verissimus* | 1 |
| | *Liphanthus barbatus* | 10 |
| | *Liphanthus longicornis* | 1 |

| | | |
|---|---|---|
| | *Liphanthus penai* | 2 |
| | *Liphanthus phasganoventris* | 1 |
| | *Liphanthus rozeni* | 2 |
| | *Liphanthus spiniventris* | 1 |
| | *Liphanthus tarsalis* | 1 |
| | *Lonchopria rufitorax* | 1 |
| | *Mourecotelles chillan* | 4 |
| | *Mourecotelles fritzi* | 3 |
| | *Mourecotelles puelche* | 1 |
| | *Mourecotelles sicheli* | 4 |
| | *Perditomorpha penai* | 2 |
| | *Spinoliella propinqua* | 1 |
| | *Spinoliella rufiventris* | 12 |
| | *Xeromelissa chillan* | 3 |
| **3** | *Calliopsis rigormortis* | 1 |
| | *Chilicola luzmarieae* | 1 |
| | *Chilicola mavida* | 1 |
| | *Chilicola randolphi* | 1 |
| | *Coelioxys wagenknechti* | 3 |
| | *Colletes flavipilosus* | 1 |
| | *Colletes guanta* | 2 |
| | *Liphanthus centralis* | 1 |
| | *Liphanthus molavi* | 1 |
| | *Trichothurgus pseudocellatus* | 1 |
| **4** | *Austropanurgus danunciae* | 1 |
| | *Caenohalictus intermedius* | 1 |
| | *Caenohalictus oblitus* | 1 |
| | *Caenohalictus pygosinuatus* | 11 |
| | *Caupolicana adusta* | 1 |
| | *Caupolicana funebris* | 1 |
| | *Chilicola colliguay* | 3 |
| | *Chilicola hahni* | 1 |
| | *Chilicola katherinae* | 1 |
| | *Chilicola lonco* | 1 |
| | *Chilicola olmue* | 2 |
| | *Chilicola orophila* | 1 |
| | *Chilicola solervicensi* | 1 |
| | *Chilicola vina* | 5 |
| | *Chilicola yali* | 1 |
| | *Coelioxys elizabeth* | 2 |
| | *Coelioxys lyprura* | 1 |

| | | |
|---|---|---|
| | *Colletes cyaniventris* | 2 |
| | *Colletes kuhlmanni* | 1 |
| | *Corynura atrovirens* | 1 |
| | *Eucerinoda gayi* | 2 |
| | *Hylaeus euxanthus* | 7 |
| | *Lasioglossum spinolae* | 14 |
| | *Liphanthus anacanthus* | 4 |
| | *Liphanthus friesellus* | 1 |
| | *Liphanthus leucostomus* | 3 |
| | *Liphanthus unifasciatus* | 2 |
| | *Lonchopria luteipes* | 1 |
| | *Megachile fasciatella* | 1 |
| | *Mourecotelles subandinus* | 1 |
| | *Notanthidium bidentatum* | 3 |
| | *Orphana inquirenda* | 6 |
| | *Parasarus spiniventris* | 1 |
| | *Perditomorpha andina* | 1 |
| | *Pseudosarus virescens* | 13 |
| | *Ruizantheda cerdai* | 2 |
| | *Spinoliella polita* | 1 |
| | *Torocolletes ibanezii* | 1 |
| | *Xeromelissa machi* | 1 |
| **5** | *Alloscirtetica vagabunda* | 4 |
| | *Austropanurgus punctatus* | 5 |
| | *Callonychium coquimbense* | 1 |
| | *Chilicola andina* | 1 |
| | *Chilicola charizard* | 1 |
| | *Chilicola diaguita* | 1 |
| | *Chilicola packeri* | 1 |
| | *Chilicola pangue* | 1 |
| | *Chilicola vicugna* | 1 |
| | *Chilimalopsis parvula* | 19 |
| | *Colletes coquimbensis* | 1 |
| | *Diadasia ruficruris* | 3 |
| | *Kelita penai* | 2 |
| | *Liphanthus domeykoi* | 1 |
| | *Liphanthus sapos* | 1 |
| | *Lonchopria heberti* | 1 |
| | *Megachile wagenknechti* | 1 |
| | *Mourecotelles atacama* | 1 |
| | *Neffapis longilingua* | 6 |

| | | |
|---|---|---|
| | *Orphana wagenknechti* | 23 |
| | *Spinoliella confusa* | 1 |
| | *Spinoliella nomadoides* | 5 |
| | *Spinoliella opaca* | 6 |
| | *Xeromelissa minuta* | 1 |
| | *Xeromelissa obscura* | 1 |
| **6** | *Chilicola erithropoda* | 1 |
| | *Chilicola guanicoe* | 1 |
| | *Chilicola travesia* | 1 |
| | *Xeromelissa brevimalaris* | 1 |
| | *Xeromelissa sielfeldi* | 1 |

Table S3 – **Indicator species per wild bee ecoregion (WBE)**

| | **Well-Sampled Assemblages** | | | | **All Unique Bee Occurrences** | | | |
|---|---|---|---|---|---|---|---|---|
| | Overview | | | | Overview | | | |
| | Total number of species: 256<br>Selected number of species: 94<br>Number of species associated to 1 group: 47<br>Number of species associated to 2 groups: 31<br>Number of species associated to 3 groups: 13<br>Number of species associated to 4 groups: 3<br>Number of species associated to 5 groups: 0 | | | | Total number of species: 423<br>Selected number of species: 132<br>Number of species associated to 1 group: 55<br>Number of species associated to 2 groups: 51<br>Number of species associated to 3 groups: 21<br>Number of species associated to 4 groups: 5<br>Number of species associated to 5 groups: 0 | | | |
| **WBE** | Species | Corr. | *P.* | Sig. | Species | Corr. | *P.* | Sig. |
| 1 | *Diphaglossa gayi* | 0.524 | 0.001 | *** | *Diphaglossa gayi* | 0.455 | 0.001 | *** |
| | *Corynura corinogaster* | 0.304 | 0.01 | ** | *Liphanthus cerdai* | 0.225 | 0.008 | ** |
| | | | | | *Callonychium chilense* | 0.199 | 0.008 | ** |
| | | | | | *Corynura ampliata* | 0.183 | 0.014 | * |
| 2 | *Corynura patagonica* | 0.548 | 0.001 | *** | *Megachile cinerea* | 0.392 | 0.001 | *** |
| | *Corynura analis* | 0.5 | 0.002 | ** | *Corynura patagonica* | 0.376 | 0.001 | *** |
| | *Parepeolus stuardi* | 0.385 | 0.047 | * | *Corynura analis* | 0.278 | 0.001 | *** |
| | *Liphanthus chillanensis* | 0.313 | 0.049 | * | *Liphanthus barbatus* | 0.227 | 0.015 | * |
| | *Liphanthus micheneri* | 0.313 | 0.049 | * | *Spinoliella rufiventris* | 0.197 | 0.017 | * |
| | *Spinoliella rufiventris* | 0.313 | 0.049 | * | *Callistochlora prothysteres* | 0.195 | 0.006 | ** |
| | | | | | *Corynura apicata* | 0.161 | 0.032 | * |
| | | | | | *Liphanthus nitidus* | 0.151 | 0.027 | * |
| 3 | *Alloscirtetica gazullai* | 0.569 | 0.002 | ** | *Anthidium espinosai* | 0.414 | 0.001 | *** |

| | | | | | | | | |
|---|---|---|---|---|---|---|---|---|
| | *Anthidium chubuti* | 0.569 | 0.002 | ** | *Anthidium chubuti* | 0.354 | 0.002 | ** |
| | *Anthidium espinosai* | 0.569 | 0.002 | ** | *Mourecotelles andinus* | 0.338 | 0.003 | ** |
| | *Caenohalictus iodurus* | 0.455 | 0.007 | ** | *Colletes fulvipes* | 0.289 | 0.001 | *** |
| | *Liphanthus andinus* | 0.455 | 0.007 | ** | *Caenohalictus iodurus* | 0.271 | 0.007 | ** |
| | *Rhophitulus evansi* | 0.455 | 0.007 | ** | *Rhophitulus evansi* | 0.271 | 0.007 | ** |
| | *Anthidium decaspilum* | 0.401 | 0.038 | * | *Alloscirtetica gazullai* | 0.23 | 0.01 | ** |
| | *Chilicola mavida* | 0.401 | 0.038 | * | *Liphanthus andinus* | 0.23 | 0.013 | * |
| | *Coelioxys wagenknechti* | 0.401 | 0.038 | * | *Megachile melanotricha* | 0.23 | 0.011 | * |
| | *Colletes guanta* | 0.401 | 0.038 | * | *Megachile distinguenda* | 0.209 | 0.008 | ** |
| | *Megachile coquimbensis* | 0.401 | 0.038 | * | *Liphanthus alicahue* | 0.153 | 0.024 | * |
| | *Mourecotelles andinus* | 0.401 | 0.038 | * | | | | |
| | *Trichothurgus pseudocellatus* | 0.401 | 0.038 | * | | | | |
| | *Megachile distinguenda* | 0.375 | 0.014 | * | | | | |
| | *Centris cineraria* | 0.303 | 0.009 | ** | | | | |
| 4 | *Anthidium chilense* | 0.292 | 0.007 | ** | *Anthidium chilense* | 0.3 | 0.001 | *** |
| | *Corynura cristata* | 0.289 | 0.025 | * | *Alloscirtetica tristrigata* | 0.209 | 0.001 | *** |
| | *Xylocopa augusti* | 0.289 | 0.019 | * | *Chalepogenus rozeni* | 0.197 | 0.003 | ** |
| | *Torocolletes fazii* | 0.259 | 0.047 | * | *Alloscirtetica gayi* | 0.186 | 0.001 | *** |
| | | | | | *Corynura cristata* | 0.186 | 0.007 | ** |
| | | | | | *Torocolletes fazii* | 0.17 | 0.012 | * |
| | | | | | *Spinolapis kalen* | 0.163 | 0.016 | * |
| | | | | | *Caupolicana hirsuta* | 0.16 | 0.012 | * |
| | | | | | *Coelioxys mapuche* | 0.152 | 0.036 | * |
| | | | | | *Liphanthus parvulus* | 0.151 | 0.02 | * |
| | | | | | *Caupolicana gayi* | 0.15 | 0.029 | * |
| | | | | | *Hylaeus punctatus* | 0.143 | 0.049 | * |
| 5 | *Neofidelia profuga* | 0.727 | 0.001 | *** | *Neofidelia profuga* | 0.497 | 0.001 | *** |
| | *Chilicolletes delahozii* | 0.535 | 0.001 | *** | *Edwyniana flavicornis* | 0.292 | 0.001 | *** |
| | *Toromelissa nemaglossa* | 0.528 | 0.001 | *** | *Chilicolletes delahozii* | 0.29 | 0.001 | *** |
| | *Orphana wagenknechti* | 0.451 | 0.004 | ** | *Perditomorpha perezi* | 0.278 | 0.001 | *** |
| | *Edwyniana flavicornis* | 0.442 | 0.001 | *** | *Orphana wagenknechti* | 0.254 | 0.001 | *** |
| | *Chilimalopsis parvula* | 0.42 | 0.004 | ** | *Penapis penai* | 0.254 | 0.001 | *** |
| | *Penapis penai* | 0.42 | 0.004 | ** | *Chilimalopsis parvula* | 0.24 | 0.001 | *** |
| | *Perditomorpha perezi* | 0.42 | 0.001 | *** | *Lonchopria similis* | 0.222 | 0.001 | *** |
| | *Kelita tuberculata* | 0.388 | 0.006 | ** | *Perditomorpha erithrogaster* | 0.22 | 0.003 | ** |
| | *Trichothurgus dubius* | 0.359 | 0.005 | ** | *Melectoides triseriatus* | 0.182 | 0.004 | ** |
| | *Epiclopus wagenknechti* | 0.353 | 0.016 | * | *Nolanomelissa toroi* | 0.178 | 0.03 | * |
| | *Chilicola rubriventris* | 0.342 | 0.013 | * | *Spinoliella opaca* | 0.178 | 0.034 | * |
| | *Perditomorpha erithrogaster* | 0.342 | 0.012 | * | *Spinoliella herbsti* | 0.171 | 0.017 | * |
| | *Lonchopria similis* | 0.336 | 0.008 | ** | *Caupolicana fulvicollis* | 0.168 | 0.012 | * |
| | *Centris rhodophthalma* | 0.309 | 0.024 | * | *Spinoliella nomadoides* | 0.159 | 0.034 | * |
| | *Kelita toroi* | 0.303 | 0.022 | * | *Chilicola vernalis* | 0.152 | 0.012 | * |
| | *Spinoliella herbsti* | 0.303 | 0.016 | * | | | | |
| 6 | *Callonychium atacamense* | 0.705 | 0.015 | * | *Centris buchholzi* | 0.354 | 0.001 | *** |
| | *Chilicola travesia* | 0.705 | 0.015 | * | *Xeromelissa rozeni* | 0.338 | 0.003 | ** |

| | | | | | | | | |
|---|---|---|---|---|---|---|---|---|
| | *Melectoides rozeni* | 0.493 | 0.032 | * | *Mesonychium garleppi* | 0.271 | 0.011 | * |
| | | | | | *Reedapis bathycyanea* | 0.223 | 0.013 | * |
| 1 + 2 | *Megachile cinerea* | 0.481 | 0.001 | *** | *Bombus dahlbomii* | 0.335 | 0.001 | *** |
| | *Coelioxys chilensis* | 0.432 | 0.003 | ** | *Ruizantheda proxima* | 0.302 | 0.001 | *** |
| | *Isepeolus lativalvis* | 0.355 | 0.008 | ** | *Bombus ruderatus* | 0.277 | 0.001 | *** |
| | *Bombus ruderatus* | 0.35 | 0.018 | * | *Chalepogenus caeruleus* | 0.254 | 0.001 | *** |
| | *Liphanthus nitidus* | 0.342 | 0.013 | * | *Cadeguala albopilosa* | 0.244 | 0.001 | *** |
| | *Callistochlora prothysteres* | 0.296 | 0.024 | * | *Coelioxys chilensis* | 0.242 | 0.002 | ** |
| | *Manuelia postica* | 0.257 | 0.045 | * | *Isepeolus lativalvis* | 0.241 | 0.002 | ** |
| | | | | | *Manuelia postica* | 0.222 | 0.001 | *** |
| | | | | | *Corynura rubella* | 0.21 | 0.004 | ** |
| | | | | | *Melissoptila dama* | 0.195 | 0.011 | * |
| | | | | | *Parepeolus stuardi* | 0.187 | 0.007 | ** |
| | | | | | *Corynura callicladura* | 0.159 | 0.043 | * |
| | | | | | *Coelioxys pergandei* | 0.132 | 0.05 | * |
| 1 + 4 | *Cadeguala occidentalis* | 0.435 | 0.001 | *** | *Bombus terrestris* | 0.213 | 0.001 | *** |
| | *Corynura herbsti* | 0.289 | 0.025 | * | *Ruizantheda mutabilis* | 0.209 | 0.003 | ** |
| | *Alloscirtetica tristrigata* | 0.277 | 0.018 | * | *Lonchopria zonalis* | 0.143 | 0.032 | * |
| | *Bombus terrestris* | 0.263 | 0.021 | * | | | | |
| 2 + 3 | *Colletes fulvipes* | 0.542 | 0.001 | *** | *Megachile santacrucensis* | 0.197 | 0.006 | ** |
| | *Megachile santacrucensis* | 0.376 | 0.016 | * | *Liphanthus chillanensis* | 0.172 | 0.025 | * |
| 2 + 4 | *Notanthidium steloides* | 0.375 | 0.001 | *** | *Colletes cyanescens* | 0.23 | 0.001 | *** |
| | *Megachile euzona* | 0.331 | 0.004 | ** | *Acamptopoeum submetallicum* | 0.181 | 0.003 | ** |
| | *Acamptopoeum submetallicum* | 0.288 | 0.014 | * | | | | |
| | *Colletes cyanescens* | 0.27 | 0.016 | * | | | | |
| 3 + 4 | - | - | - | - | *Xylocopa augusti* | 0.215 | 0.002 | ** |
| | | | | | *Centris nigerrima* | 0.206 | 0.001 | *** |
| | | | | | *Corynura herbsti* | 0.206 | 0.003 | ** |
| | | | | | *Caupolicana quadrifasciata* | 0.16 | 0.005 | ** |
| | | | | | *Anthidium manicatum* | 0.15 | 0.025 | * |
| | | | | | *Epiclopus lendlianus* | 0.138 | 0.048 | * |
| 3 + 5 | *Alloscirtetica rufitarsis* | 0.415 | 0.002 | ** | *Alloscirtetica rufitarsis* | 0.238 | 0.001 | *** |
| | *Liphanthus coquimbensis* | 0.362 | 0.01 | ** | *Liphanthus coquimbensis* | 0.215 | 0.002 | ** |
| | *Mourecotelles incahuasi* | 0.301 | 0.026 | * | *Mourecotelles incahuasi* | 0.199 | 0.013 | * |
| | *Mourecotelles ruizii* | 0.296 | 0.016 | * | *Mourecotelles ruizii* | 0.193 | 0.005 | ** |
| | | | | | *Caenohalictus rostraticeps* | 0.171 | 0.022 | * |
| | | | | | *Chilicola rubriventris* | 0.17 | 0.01 | ** |
| | | | | | *Megachile rancaguensis* | 0.146 | 0.033 | * |
| 3 + 6 | *Liphanthus alicahue* | 0.43 | 0.001 | *** | *Alloscirtetica porteri* | 0.211 | 0.011 | * |
| 4 + 5 | *Diadasia chilensis* | 0.339 | 0.001 | *** | *Trichothurgus dubius* | 0.167 | 0.015 | * |
| 5 + 6 | *Colletes atripes* | 0.61 | 0.001 | *** | *Parasarus atacamensis* | 0.452 | 0.001 | *** |
| | *Parasarus atacamensis* | 0.61 | 0.001 | *** | *Colletes atripes* | 0.445 | 0.001 | *** |
| | *Perditomorpha atacama* | 0.433 | 0.004 | ** | *Perditomorpha atacama* | 0.372 | 0.001 | *** |
| | *Edwyniana herrerae* | 0.42 | 0.002 | ** | *Edwyniana herrerae* | 0.327 | 0.001 | *** |
| | *Perditomorpha flavitarsus* | 0.339 | 0.023 | * | *Epiclopus wagenknechti* | 0.322 | 0.001 | *** |

| | | | | | | | | |
|---|---|---|---|---|---|---|---|---|
| | *Spinoliella rozeni* | 0.339 | 0.027 | * | *Toromelissa nemaglossa* | 0.317 | 0.001 | *** |
| | *Centris chilensis* | 0.266 | 0.015 | * | *Centris chilensis* | 0.292 | 0.001 | *** |
| | | | | | *Penapis moldenkei* | 0.275 | 0.001 | *** |
| | | | | | *Alloscirtetica chilena* | 0.265 | 0.001 | *** |
| | | | | | *Perditomorpha flavitarsus* | 0.243 | 0.001 | *** |
| | | | | | *Isepeolus wagenknechti* | 0.236 | 0.002 | ** |
| | | | | | *Kelita tuberculata* | 0.236 | 0.001 | *** |
| | | | | | *Spinoliella rozeni* | 0.207 | 0.01 | ** |
| | | | | | *Neofidelia longirostris* | 0.193 | 0.002 | ** |
| | | | | | *Kelita toroi* | 0.181 | 0.012 | * |
| | | | | | *Centris rhodophthalma* | 0.179 | 0.007 | ** |
| 1 + 2 + 3 | *Megachile semirufa* | 0.554 | 0.001 | *** | *Megachile semirufa* | 0.267 | 0.001 | *** |
| | *Chalepogenus caeruleus* | 0.342 | 0.017 | * | | | | |
| 1 + 2 + 4 | *Ruizantheda proxima* | 0.419 | 0.001 | *** | *Cadeguala occidentalis* | 0.306 | 0.001 | *** |
| | *Svastrides melanura* | 0.377 | 0.002 | ** | *Manuelia gayatina* | 0.246 | 0.001 | *** |
| | *Manuelia gayatina* | 0.363 | 0.001 | *** | *Megachile pollinosa* | 0.219 | 0.001 | *** |
| | *Cadeguala albopilosa* | 0.347 | 0.004 | ** | *Manuelia gayi* | 0.208 | 0.003 | ** |
| | *Manuelia gayi* | 0.301 | 0.006 | ** | *Doeringiella gayi* | 0.15 | 0.03 | * |
| | *Anthidium gayi* | 0.295 | 0.011 | * | | | | |
| 1 + 3 + 4 | *Ruizantheda mutabilis* | 0.321 | 0.004 | ** | *Sphecodes chilensis* | 0.167 | 0.006 | ** |
| | *Isepeolus septemnotatus* | 0.281 | 0.035 | * | *Isepeolus septemnotatus* | 0.166 | 0.009 | ** |
| 2 + 3 + 4 | *Calliopsis trifasciata* | 0.493 | 0.001 | *** | *Calliopsis trifasciata* | 0.294 | 0.001 | *** |
| | | | | | *Notanthidium steloides* | 0.281 | 0.001 | *** |
| | | | | | *Corynura chilensis* | 0.23 | 0.001 | *** |
| | | | | | *Megachile euzona* | 0.213 | 0.001 | *** |
| | | | | | *Centris cineraria* | 0.2 | 0.001 | *** |
| | | | | | *Callistochlora chloris* | 0.171 | 0.005 | ** |
| 3 + 4 + 5 | *Megachile saulcyi* | 0.369 | 0.001 | *** | *Diadasia chilensis* | 0.307 | 0.001 | *** |
| | *Liphanthus sabulosus* | 0.292 | 0.008 | ** | *Liphanthus sabulosus* | 0.216 | 0.002 | ** |
| | | | | | *Alloscirtetica valparadisaea* | 0.177 | 0.001 | *** |
| | | | | | *Megachile rotundata* | 0.177 | 0.002 | ** |
| | | | | | *Reedapis semicyanea* | 0.156 | 0.011 | * |
| | | | | | *Mourecotelles mixtus* | 0.14 | 0.05 | * |
| 3 + 5 + 6 | - | - | - | - | *Colletes sulcatus* | 0.179 | 0.005 | ** |
| 1 + 2 + 3 + 4 | *Bombus dahlbomii* | 0.395 | 0.001 | *** | *Svastrides melanura* | 0.295 | 0.001 | *** |
| | *Corynura chilensis* | 0.311 | 0.008 | ** | *Colletes nigritulus* | 0.184 | 0.005 | ** |
| | | | | | *Anthidium gayi* | 0.177 | 0.005 | ** |
| | | | | | *Corynura corinogaster* | 0.151 | 0.023 | * |
| 1 + 2 + 4 + 6 | *Megachile pollinosa* | 0.382 | 0.001 | *** | - | - | - | - |
| 3 + 4 + 5 + 6 | | | | | *Megachile saulcyi* | 0.285 | 0.001 | *** |

**Supplementary Information**

At present, over 480 species have been described for Chile (Ascher and Pickering, 2020), and many well curated morphospecies await description and more remain to be collected and recognized as additional areas, underrepresented in current collections, are surveyed and integrative approaches to taxonomy are applied (Packer & Ruz 2016). A handful of researchers and research groups have produced most species descriptions, starting with Herbst and expanding in the 70s and 80s with the work of Toro and his collaborators (*e.g.,* Rojas and Toro 2000, Ruz and Toro 1983, Toro and Rodríguez 1998, Toro and Moldenke 1979, Toro and Ruz 1969). More recently, contributions have been made through classical taxonomy as well as modern integrative taxonomy approaches (*e.g.,* Ferrari 2017, González-Vaquero & Galvani 2016, Monckton 2016, Packer & Dumesh 2014, Roig-Alsina 1991, Roig-Alsina 1999, Vivallo 2009, Vivallo 2014, Vivallo et al., 2003, Vivallo 2013; Henríquez-Piskulich et al., 2020). In addition to continued work on the fauna of Central Chile, work by Packer et al., (e.g. Dumesh & Packer 2013; Packer & Graham, 2020), have documented a rich bee fauna in Northern Chile (north of the region treated here, i.e. in Arica and Parinacota that was less well surveyed by historical workers).

**Supplementary Results**

- The five most common species (most unique records in the dataset of temporally and spatially unique occurrences) were *Acamptopoeum submetallicum* (Spinola, 1851) (Hymenoptera: Andrenidae) (376), *Anthidium chilense* Spinola, 1851 (Hymenoptera: Megachilidae) (369), *Megachile saulcyi* Guérin-Méneville, 1845 (Hymenoptera: Megachilidae) (368), *Bombus dahlbomii* (358), and *Centris chilensis* (Spinola, 1851) (Hymenoptera: Apidae).

- The five most common species in the well-sampled grid cells/sites were *Anthidium chilense* (with 89 grid cells), *Megachile saulcyi* (89), *Acamptopoeum submetallicum* (87), *Colletes cyanescens* (Haliday, 1836) (Hymenoptera: Colletidae) (75), and *Megachile pollinosa* Spinola, 1851 (Hymenoptera: Megachilidae) (71).
